# Signal peptidase SpsB coordinates staphylococcal cell cycle, surface protein septal trafficking and LTA synthesis

**DOI:** 10.1101/2024.08.20.608893

**Authors:** Ran Zhang, Yaosheng Jia, Salvatore J. Scaffidi, Jesper J. Madsen, Wenqi Yu

**Affiliations:** Department of Molecular Biosciences, College of Arts and Sciences, University of South Florida, Tampa, Florida 33620, United States of America; Department of Molecular Medicine, Morsani College of Medicine; Center for Global Health and Infectious Diseases Research, Global and Planetary Health, College of Public Health, University of South Florida, Tampa, Florida 33620, United States of America

**Keywords:** SpsB, YSIRK/G-S signal peptide, LtaS, staphylococcal cell cycle, staphylococcal protein A (SpA)

## Abstract

Many cell wall anchored surface proteins of Gram-positive bacteria harbor a highly conserved YSIRK/G-S signal peptide (SP_YSIRK+_), which deposits surface protein precursors at the cell division septum where they are subsequently anchored to septal peptidoglycan. Previously we identified that LtaS-mediated lipoteichoic acid (LTA) synthesis regulates septal trafficking of YSIRK+ proteins in *S. aureus*. Interestingly, both LtaS and SP_YSIRK+_ are cleaved by the signal peptidase SpsB, but the biological implications remain unclear. Here we show that SpsB is required for cleaving SP_SpA(YSIRK+)_ of staphylococcal surface protein A (SpA). Depletion of *spsB* not only diminished SP_SpA_ processing but also abolished SpA septal localization. The mis-localization is attributed to the cleavage activity of SpsB, as an A37P mutation of SP_SpA_ that disrupted SpsB cleavage also abrogated SpA septal localization. Strikingly, depletion of *spsB* led to aberrant cell morphology, cell cycle arrest and daughter cell separation defects. Localization studies showed that SpsB predominantly localized at the septum of dividing staphylococcal cells. Finally, we show that SpsB spatially regulates LtaS as *spsB* depletion enriched LtaS at the septum. Collectively, the data suggest a new dual-mechanism model mediated by SpsB: the abundant YSIRK+ proteins are efficiently processed by septal localized SpsB; SpsB cleaves LtaS at the septum, which spatially regulates LtaS activity contributing to YSIRK+ proteins septal trafficking. The study identifies SpsB as a novel and key regulator orchestrating protein secretion, cell cycle and cell envelope biogenesis.

**Importance:** Surface proteins containing a YSIRK/G-S positive signal peptide are widely distributed in Gram-positive bacteria and play essential roles in bacterial pathogenesis. They are highly expressed proteins that are enriched at the septum during cell division. The biogenesis of these proteins is coordinated with cell cycle and LTA synthesis. The current study identified the staphylococcal signal peptidase SpsB as a key determinant in regulating surface protein septal trafficking. Furthermore, this study highlights the novel functions of SpsB in coordinating LtaS-mediated LTA production and regulating staphylococcal cell cycle. As SpsB, YSIRK+ proteins and LTA synthesis are widely distributed and conserved, the mechanisms identified here may be shared across Gram-positive bacteria.

## Introduction

The cell wall anchored surface proteins of Gram-positive bacteria are widely distributed and constitute an integral part of bacterial cell envelope. These proteins are covalently attached to cell wall peptidoglycan and displayed on the bacterial cell surface, which are essential in bacterial interactions with the environment (Marraffini *et al*, 2006). In the Gram-positive pathogen *Staphylococcus aureus* for example (Tong *et al*, 2015), cell wall anchored surface proteins play vital roles in adhesion, biofilm formation, nutrient acquisition, antibiotics resistance and immune evasion (Foster *et al*, 2014; Schneewind & Missiakas, 2019).

The biochemical pathway of surface protein trafficking (the sorting pathway) is conserved in most Gram-positive bacteria and best understood in *S. aureus.* In the sorting pathway, surface protein precursors with an N-terminal signal peptide (SP) and a C-terminal cell wall sorting signal are produced in the cytoplasm (Schneewind *et al*, 1992). The SP mediates preprotein membrane translocation via the Sec secretion pathway (Yu *et al*, 2018). Upon membrane translocation through the SecYEG translocon, the SP is cleaved by the type I signal peptidase, SpsB in *S. aureus* (Madsen & Yu, 2024). Subsequently, the membrane bound transpeptidase sortase A (SrtA) covalently attaches surface protein precursors to the cell wall precursor lipid II, which is further incorporated to the mature cell wall meshwork during cell wall biosynthesis (Perry *et al*, 2002; Ton-That *et al*, 1997).

Intriguingly, many surface proteins of Gram-positive bacteria contain a specific N-terminal SP with a highly conserved YSIRK/G-S motif (abbreviated as SP_YSIRK+_) (Rosenstein & Götz, 2000; Tettelin *et al*, 2001). The YSIRK/G-S motif has a conserved pattern of YSIRKxxxGxxS positioned at the beginning of the hydrophobic region of the SP. Signal peptides containing YSIRK/G-S motif have been shown to target proteins to the cell division septum (Carlsson *et al*, 2006; DeDent *et al*, 2008; Yu & Götz, 2012). The septum and septal peptidoglycan (cross-wall) compartment constitute the center of cell division and cell envelope assembly. The proposed paradigm suggests that SP_YSIRK+_ directs protein secretion at the septum; septal secreted proteins are subsequently anchored to the cross-wall (Carlsson *et al*., 2006; DeDent *et al*., 2008); upon cell separation, cross-wall anchored surface proteins are displayed on the new hemisphere of the daughter cells. However, the underlying mechanisms are still largely unknown.

In our previous studies, we investigated the septal trafficking of staphylococcal protein A (SpA), as an archetype of YSIRK/G-S proteins. SpA is one of the most abundant proteins in *S. aureus* and a well-known virulence factor for its function of binding host immunoglobulin (Falugi *et al*, 2013; Forsgren & Sjöquist, 1966; Kim *et al*, 2016). We have shown that SpA precursor interacts with SecA and engages SecA-dependent Sec secretion pathway. SecA is required for SpA secretion but does not determine SpA septal localization, as SecA localizes circumferentially in proximity of cytoplasmic membrane (Yu *et al*., 2018). Furthermore, we found that LtaS-mediated lipoteichoic acid (LTA) synthesis spatially regulates SpA biogenesis (Yu *et al*., 2018; Zhang *et al*, 2021). The activity of LtaS is required for SpA septal trafficking (Ibrahim *et al*, 2024; Zhang *et al*., 2021). LtaS has been shown to localize at the septum (Reichmann *et al*, 2014), whereas LTA is predominantly found at the cell periphery (Zhang *et al*., 2021). Interestingly, while LtaS is not a typical secreted protein, it is cleaved by SpsB between its extracellular enzymatic domain (eLtaS) and transmembrane domains, resulting in inactivation of LtaS (Wormann *et al*, 2011). A recent study suggests that the cleavage slowly releases eLtaS, which temporally regulates LtaS activity and YSIRK+ protein cross-wall targeting (Ibrahim *et al*., 2024).

As SpsB cleaves both signal peptides and LtaS, we aimed to elucidate the functions of SpsB in this study. Our results show that SpsB-mediated SP cleavage is required for SP_SpA(YSIRK+)_-mediated SpA septal localization. SpsB was predominantly found at the septum and depletion of *spsB* leads to strong cell cycle arrest and cell separation defects. Moreover, depletion of *spsB* enriched LtaS localization at the septum. Thus, our study uncovers a novel function of SpsB as a cell cycle regulator and defines its key role in spatiotemporal regulation of YSIRK proteins secretion and LtaS-mediated LTA synthesis.

## Results

### Depletion of *spsB* abolished septal localization of SP_SpA(YSIRK+)_-SpA* but does not affect the peripheral localization of SP_SplE(YSIRK-)_-SpA*

A *spsB* depletion strain ANG2009 (referred to as SEJ1*ispsB* here) was constructed previously (Wormann *et al*., 2011). In this strain, native *spsB* is deleted and a single copy of *spsB* is placed under P*_spac_* promoter at an ectopic locus (**Fig. 1A**). The expression of *spsB* is depleted in the absence of IPTG and induced upon IPTG addition. The parental strain SEJ1 is a *spa* marker-less deletion mutant of *S. aureus* RN4220 (referred to as SEJ1 WT here). We deleted the *srtA* gene in *spsB* depletion mutant generating *srtA/ispsB* double mutant. We showed previously that full-length SpA contains an additional cross-wall targeting LysM domain (Zhang *et al*., 2021). To eliminate any potential interference by the LysM domain, here we used SpA_ΔLysM_ as our reporter protein whose cross-wall localization is solely mediated by its signal peptide. For simplicity, SpA_ΔLysM_ is designated as SpA* in this report. We constructed pKK30*itet-spa*\*, whereby SpA* is expressed under anhydro-tetracycline (ATc)-inducible P*_tet_* promoter (**Fig. 1A**). The reporter plasmid, together with its empty vector, were transformed to SEJ1 WT, *ispsB*, *srtA* single and double mutants. Bacterial cultures were grown with IPTG overnight, washed and refreshed in media with and without IPTG and with ATc for 3 hours to analyze the phenotypes.

**Fig. 1.**
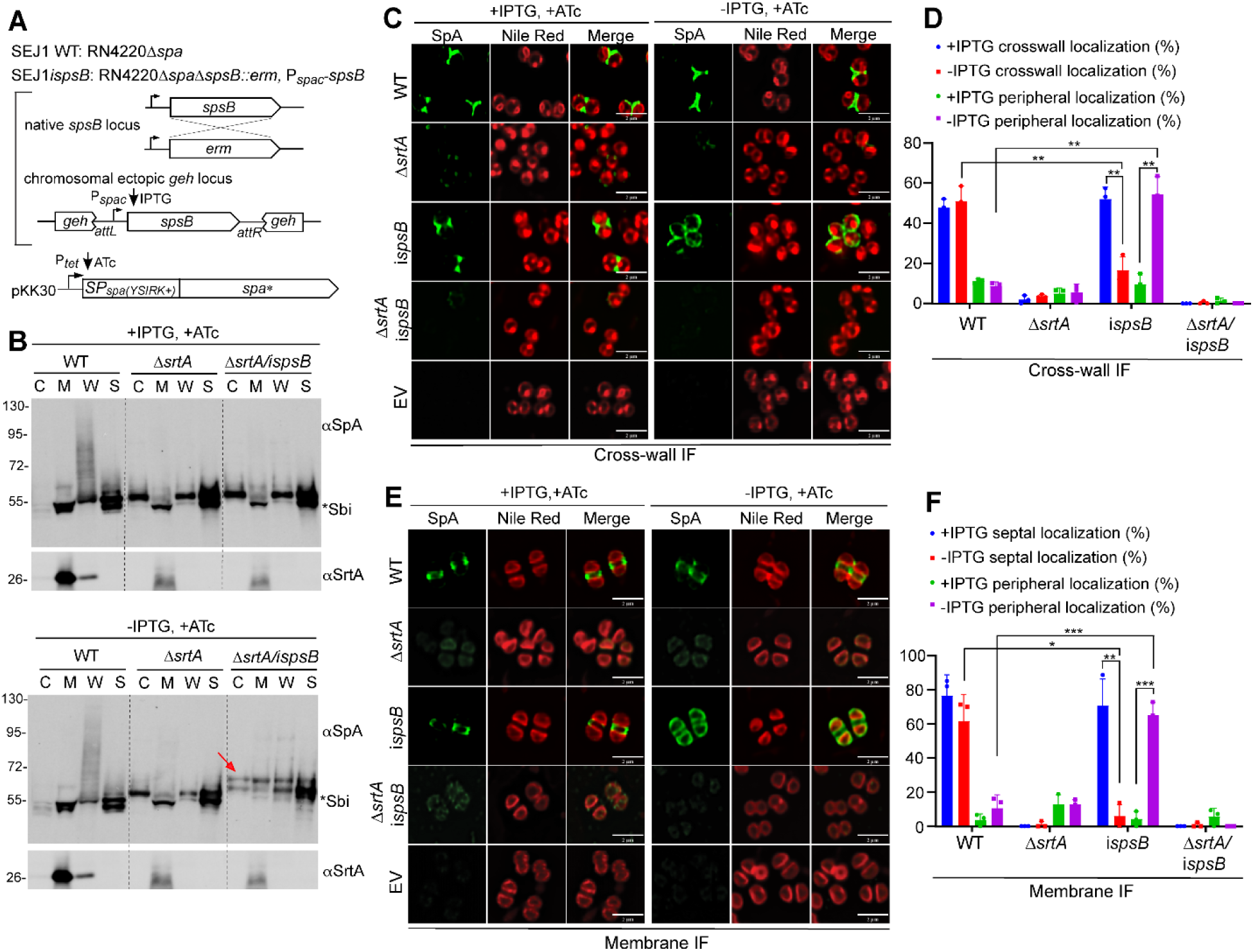
*spsB* depletion abolished septal localization of SP_SpA(YSIRK+)_-SpA*. **(A)** Genetic background of the strains. All mutants were constructed in SEJ1 (RN4220Δ*spa*) referred to as WT. In the *spsB* depletion mutant (i*spsB*), the native *spsB* was deleted and a single copy of *spsB* is expressed under IPTG-inducible P*_spac_* at an ectopic *geh* locus (Wormann *et al*., 2011). Plasmid-borne SpA* (SpA lacking the LysM domain) is expressed under ATc-inducible promotor in SEJ1 WT, Δ*srtA* and i*spsB* single and double mutants. Bacterial cultures were grown with IPTG (*spsB*-induced) and without IPTG (*spsB*-depleted) and with ATc (to induce *spa*\*) for 3 hours and subjected to experiments in B-F. **(B)** Cell fractionation and immunoblot analysis of SpA*. C, cytoplasm; M, cell membrane; W, cell wall; S, supernatant. The red arrow indicates unprocessed SP-bearing precursors. Asterisk indicates non-specific Sbi bands. Sortase A (αSrtA) blot serves as a loading and fractionation control. Numbers on the left indicate protein ladder in kDa. **(C)** Localization of newly synthesized SpA* on staphylococcal cell surface revealed by cross-wall immunofluorescence (IF) microscopy. Nile red stains cell membrane. Scale bar, 2 µm. **(D)** Quantification of SpA* cross-wall and peripheral localization from the images represented in panel C. **(E)** SpA* localization on protoplasts revealed by membrane IF. **(F)** Quantification of SpA* septal and peripheral localization from the images represented in panel E. Representative images and quantification are from three independent experiments. Unpaired t-test with Welch’s correction was performed for statistical analysis: *p <0.05; **p <0.005; ***p <0.0005; ****p <0.0001.

To examine whether *spsB* depletion impaired SP_SpA_ cleavage, we performed cell fractionation and immunoblotting. Staphylococcal cell cultures were separated into cytoplasm (C), membrane (M), cell wall (W) and supernatant (S) fractions and immunoblotted with SpA antibody. Consistent with previous studies, SpA* displayed smear-like pattern in the cell wall fraction with some proteins released to the supernatant in SEJ1 WT cells (**Fig. 1B**, WT, −/+IPTG) (Zhang *et al*., 2021). As expected, SpA* accumulated in the cytoplasm and was released to the supernatant in Δ*srtA*, as SpA* cannot be anchored to the cell wall without SrtA (**Fig. 1B**, Δ*srtA, −/+*IPTG). The Δ*srtA/ispsB* mutant showed the same immunoblot pattern as Δ*srtA* in the presence of IPTG (*spsB*-induced). In the absence of IPTG (*spsB*-depleted), a slow migrating protein band was found in the cytoplasm, membrane and cell wall fractions of Δ*srtA/ispsB* (**Fig. 1B**, red arrow, Δ*srtA/ispsB,* -IPTG). The slow migrating band represents SP-bearing precursors that were not processed by SpsB. It is known that alternative cleavage occurs when SpsB is inhibited or depleted (Wormann *et al*., 2011). A second band that ran underneath the dominant slow-migrating band likely resulted from alternative partial processing (**Fig. 1B**). *S. aureus* contains another non-specific IgG binding protein Sbi, which is a secreted protein that can be found in the membrane and supernatant fractions (Smith *et al*, 2012; Zhang *et al*., 2021). As Sbi is only slightly smaller (theoretical mass 47 kD) than SpA* (theoretical mass 50 kD), we compared samples with the empty vector control to distinguish Sbi and SpA* precursors (**Fig. S1**). In all strains tested, the slowly migrating SpA* bands were found upon *spsB* depletion (red arrows, *ispsB* or Δ*srtA/ispsB,*), which evidently migrated above the Sbi bands. In conclusion, *spsB* depletion is functional, which impairs the cleavage of SP_SpA_.

To examine SpA* and its precursor localization, we used two immunofluorescence (IF) microscopy methods that we previously established: cross-wall IF and membrane IF (Scaffidi *et al*, 2021; Scaffidi & Yu, 2024; Yu *et al*., 2018). In the cross-wall IF experiment, the pre-existing proteins on the cell surface are removed by trypsin and cells are grown again in fresh medium containing trypsin inhibitor for 20 min (equivalent to roughly one round of cell cycle) to allow protein regeneration. Cells are then fixed, stained with primary and secondary antibodies and subjected to microscopy analysis. In short, cross-wall IF reveals the deposition of newly emerged surface proteins on the bacterial cell surface. In the membrane IF experiment, trypsinized cells are fixed and digested with staphylococcal cell wall hydrolase lysostaphin to remove the cell wall peptidoglycan; the resulting protoplasts are fixed and stained with primary and secondary antibodies for microscopy analysis. We routinely use fluorescent vancomycin (Van-FL), a cell wall-binding dye, to confirm that the majority of peptidoglycan is removed while some weak Van-FL signals may remain at the septum (Scaffidi *et al*., 2021; Zhang *et al*., 2021). Importantly, lysostaphin digestion separates two daughter cells, which allows antibody penetration and detection at the septal membrane (Fig. 1E compared to Fig. 1C, for example) (Scaffidi *et al*., 2021; Yu *et al*., 2018). In short, the membrane IF reveals protein localization underneath peptidoglycan at the membrane-proximal compartment including septum.

Results from cross-wall IF showed that SpA* localized at the cross-wall in WT cells but mis-localized to the cell periphery in *spsB*-depleted cells. (**Fig. 1C**, *ispsB*, -IPTG, +ATc). Adding IPTG restored SpA* cross-wall localization. The SpA* signal was barely detectable in the Δ*srtA* single and Δ*srtA/ispsB* double mutant. This is expected as SrtA-mediated anchoring is required to display proteins on the bacterial cell surface. Similar localization patterns were found from the membrane IF experiment (**Fig. 1E**). SpA* dispersed all over the cell membrane when *spsB* was depleted, in contrast to its septal localization in WT cells or when *spsB* was induced with IPTG (**Fig. 1E**). Surprisingly, SpA* signals were still very weak in the Δ*srtA/ispsB* double mutant (Fig. 1E, Δ*srtA/ispsB,* -IPTG, +ATc), as immunoblotting showed accumulation of SpA* precursors in the membrane fraction. Presumably, the amount detected by immunoblot was not enough to be captured by IF. Quantification of microscopy images clearly indicates that *spsB* depletion abolished SpA* cross-wall and septal localization (**Fig. 1DF**).

To examine whether SpsB affects non-YSIRK signal peptide processing and precursor localization, we performed the same set of experiments with SP_SplE_-SpA* fusion (**Fig. 2A**). SplE is a secreted serine proteinase of *S. aureus* (Gimza *et al*, 2019; Reed *et al*, 2001). We used SP_SplE_ as a non-YSIRK signal peptide representative because it has the same length as SP_SpA_ (36 a.a.). Cross-wall IF and membrane IF experiments showed that SP_SplE_-SpA* distributed all over the cells; *spsB* depletion had no obvious effect on the peripheral localization of SP_SplE_-SpA* (**Fig. 2CE**). Similar to SP_SpA_-SpA*, SP_SplE_-bearing precursors accumulated in the cytoplasm, membrane and cell wall fractions upon *spsB* depletion (**Fig. 2B, Fig. S1**). These results indicate that while SpsB also cleaves SP_SplE_, depletion of *spsB* does not affect the localization of SP_SplE_-SpA*.

**Fig. 2.**
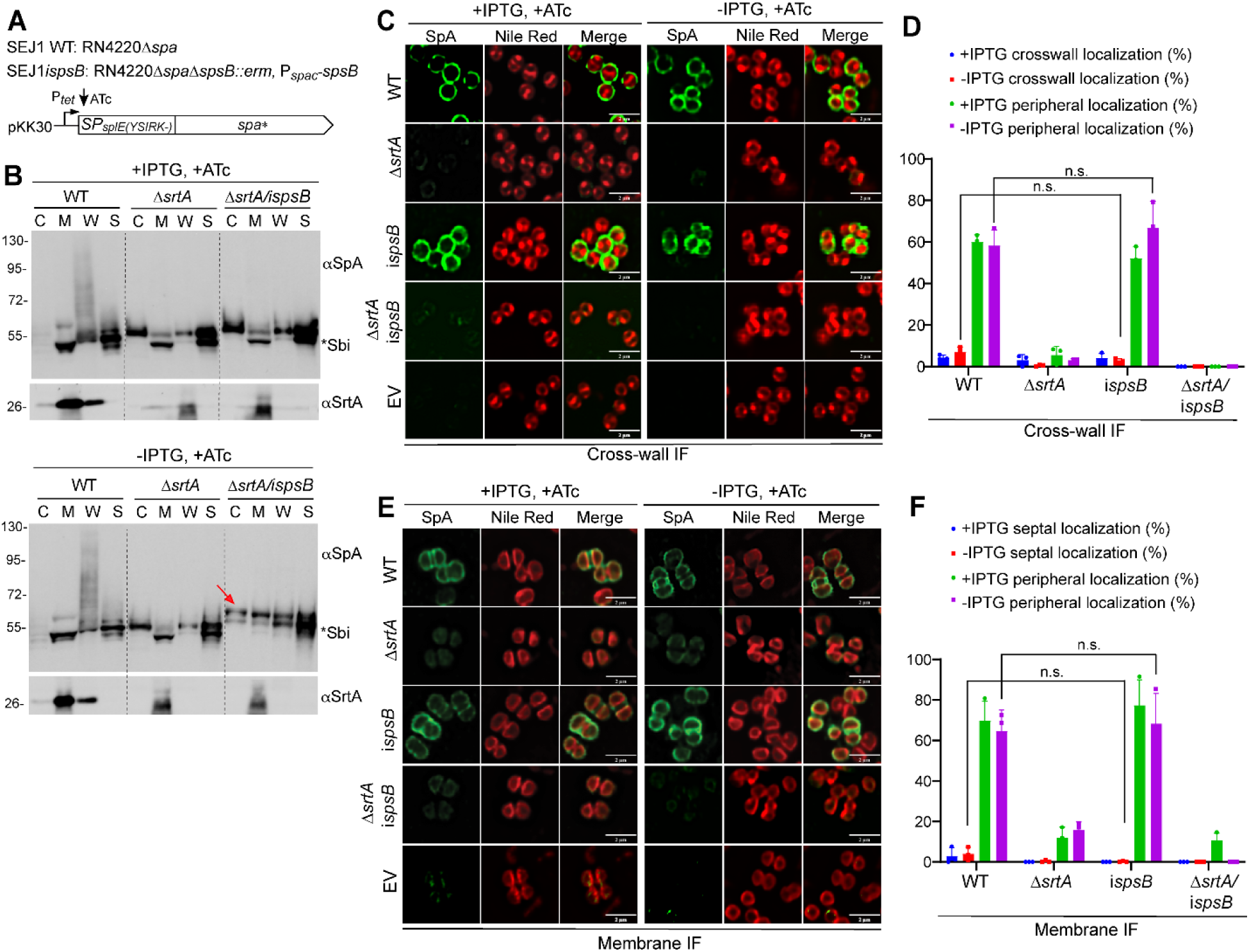
*spsB* depletion did not affect the peripheral localization of SP_SplE(YSIRK-)_-SpA*. **(A)** Genetic background of the strains. All the strains are the same as those in Fig. 1 except that SP_SplE(YSIRK-)_ replaced SP_SpA(YSIRK+)_ to fuse with SpA*. **(B)** Cell fractionation and immunoblot analysis of SpA* in the cytoplasm (C), cell membrane (M), cell wall (W), and the supernatant (S). The red arrow indicates unprocessed SP-bearing precursors. The asterisk indicates non-specific Sbi bands. The αSrtA blot is a loading and fractionation control. Numbers on the left indicate protein ladder in kDa. **(C)** Localization of newly synthesized SpA* on staphylococcal cell surface revealed by cross-wall IF. Nile red stains cell membrane. Scale bar, 2 µm. **(D)** Quantification of SpA* cross-wall and peripheral localization from the images represented in panel C. **(E)** SpA* localization on protoplasts revealed by membrane IF. **(F)** Quantification of SpA* septal and peripheral localization from the images represented in panel E. Representative images and quantification are from three independent experiments. Unpaired t-test with Welch’s correction was performed for statistical analysis: *p <0.05; **p <0.005; ***p <0.0005; ****p <0.0001.

### Efficient signal peptide cleavage by SpsB is required for SP_SpA(YSIRK+)_-SpA* septal trafficking

The above results showed that SpsB is required for SP_SpA_-SpA* septal localization. We then asked whether the phenotype was dependent on SpsB-mediated cleavage. To test this, we generated a point mutant of A37P in SP_SpA_ to disrupt SpsB-mediated cleavage. SP_SpA_ contains a typical SpsB cleavage site ‘AXA_36_’ at the C-terminal end (**Fig. 3A**). Amino acid mutation at the P1’-position right after the AXA motif is known to abrogate signal peptidase cleavage (Barkocy-Gallagher & Bassford, 1992). Indeed, cell fractionation and immunoblotting showed that the A37P variant accumulated slow-migrating SP-bearing precursors in the cytoplasm and membrane fractions of WT and *ΔsrtA* cells (**Fig. 3F**, red arrow). The accumulation is more dominant in *ΔsrtA* than WT. This is likely because unprocessed SP_SpA_A37P_-bearing precursors can be anchored to the cell wall in WT cells, in a manner similar to mobilizing a lipoprotein to the cell wall (Navarre *et al*, 1996). Nevertheless, the apparent precursor accumulation indicated that A37P point mutation disrupted SpsB-mediated cleavage.

**Fig. 3.**
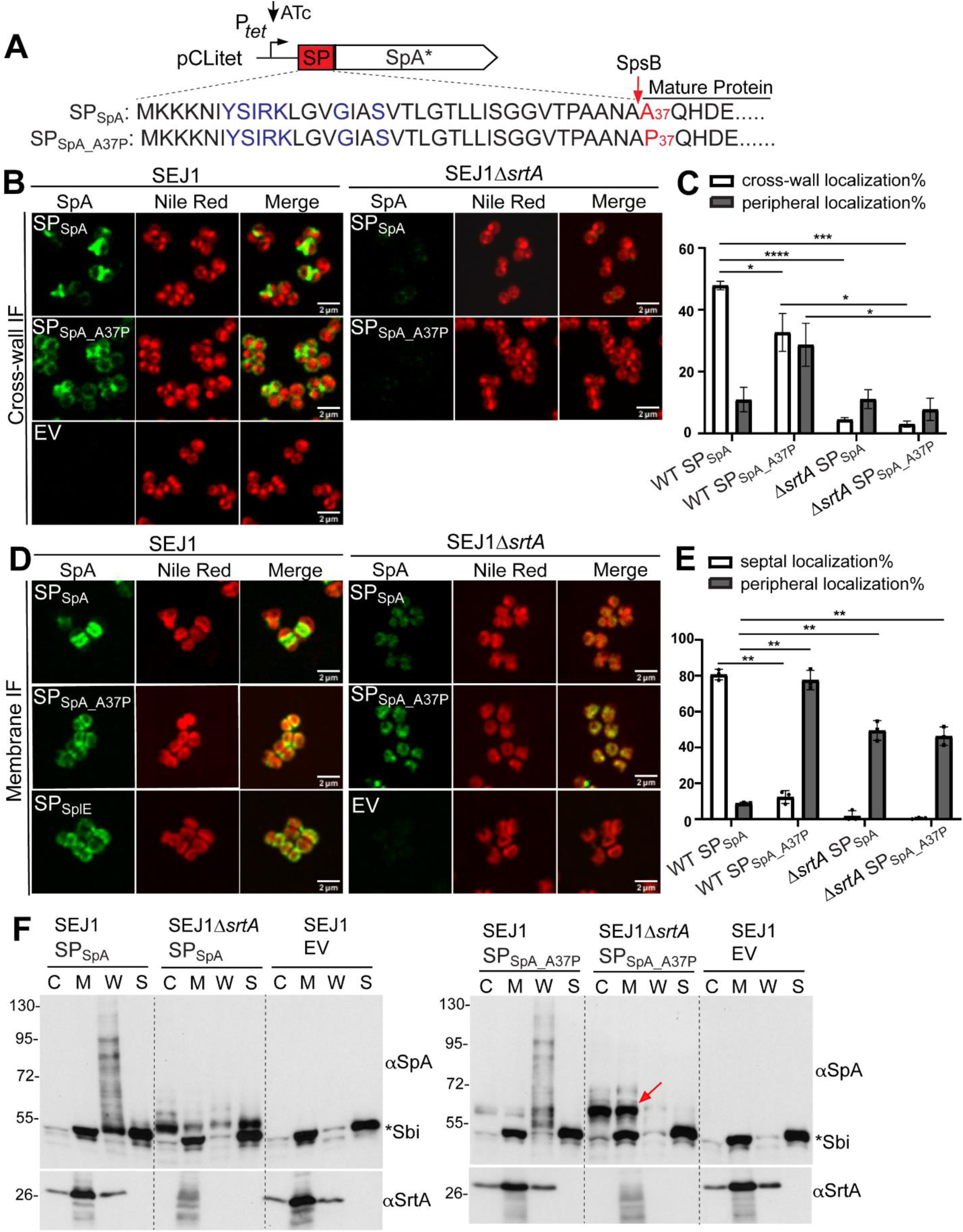
Efficient signal peptide cleavage by SpsB is required for SP_SpA(YSIRK+)_-SpA* septal trafficking. **(A)** Illustration of A37P amino acid mutation in SP_SpA_. The YSIRK/G-S motif is marked in blue. **(B)** Localization of newly synthesized SpA* on staphylococcal cell surface revealed by cross-wall IF. Nile red stains cell membrane. Scale bar, 2 µm. **(C)** Quantification of SpA* cross-wall and peripheral localization from the images represented in panel B. **(D)** SpA* localization on protoplasts revealed by membrane IF. **(E)** Quantification of SpA* septal and peripheral localization from the images represented in panel D. **(F)** Cell fractionation and immunoblot analysis of SpA* in the cytoplasm (C), cell membrane (M), cell wall (W), and the supernatant (S). The red arrow indicates unprocessed SP-bearing precursors. The asterisk indicates non-specific Sbi bands. The αSrtA blot is a loading and fractionation control. Numbers on the left indicate protein ladder in kDa. Representative images and quantification are from three independent experiments. Unpaired t-test with Welch’s correction was performed for statistical analysis: *p <0.05; **p <0.005; ***p <0.0005; ****p <0.0001.

From the cross-wall IF experiment, SP_SpA_A37P_ showed diminished cross-wall and increased peripheral wall localization in WT cells (**Fig. 3BC**). Barely any signals were detected in the *ΔsrtA* mutant as expected. Similar results were obtained from a membrane IF experiment (**Fig. 3DE**). SP_SpA_A37P_ localized circumferentially in the WT cells in contrast to septal localization of SP_SpA_. We were able to detect weak and diffused signals of SP_SpA_A37P_ all over the cell membrane in *ΔsrtA*, which likely resulted from accumulation of unprocessed SP-bearing precursors in the membrane fraction (**Fig. 3F**, red arrow). Based on these data, we concluded that signal peptide cleavage is required to deposit SpA* at the septum, which are subsequently anchored to the cross-wall.

### Depletion of *spsB* led to cell cycle arrest, cell separation and cell wall synthesis defects

While working with the *spsB* depletion mutant, we were intrigued by its aberrant cell morphology. Bacterial cultures were grown without IPTG for 3 hours and then diluted again in fresh media without IPTG for another 3 hours (6 hours of depletion) to further deplete *spsB* **(Fig. 4D)**. As expected, *spsB* depletion (*ispsB* and Δ*srtA/ispsB*) triggered severe growth defect as *spsB* is an essential gene (**Fig. 4D**). When analyzed under the microscope, *spsB*-depleted cells displayed cell cycle arrest and cell separation defects (**Fig. 4A-C, Fig. S2**). Cells showed heterogenous cell sizes, clustered together with irregular and multiple septa formation. Quantification of cell cycle progression showed that a significant number of *spsB*-depleted cells remained in phase 3, at which septa were formed but cells were unable to separate (**Fig. 4B**). As a result of cell separation defects, cells formed tetrads or larger clusters which were quantified in Fig. 4C (**Fig. 4C**). The aberrant cell morphology upon *spsB* depletion was more severe after 6 hours of depletion (**Fig. S2**). These distinct morphological defects suggest a key role of SpsB in regulating staphylococcal cell cycle.

**Fig. 4.**
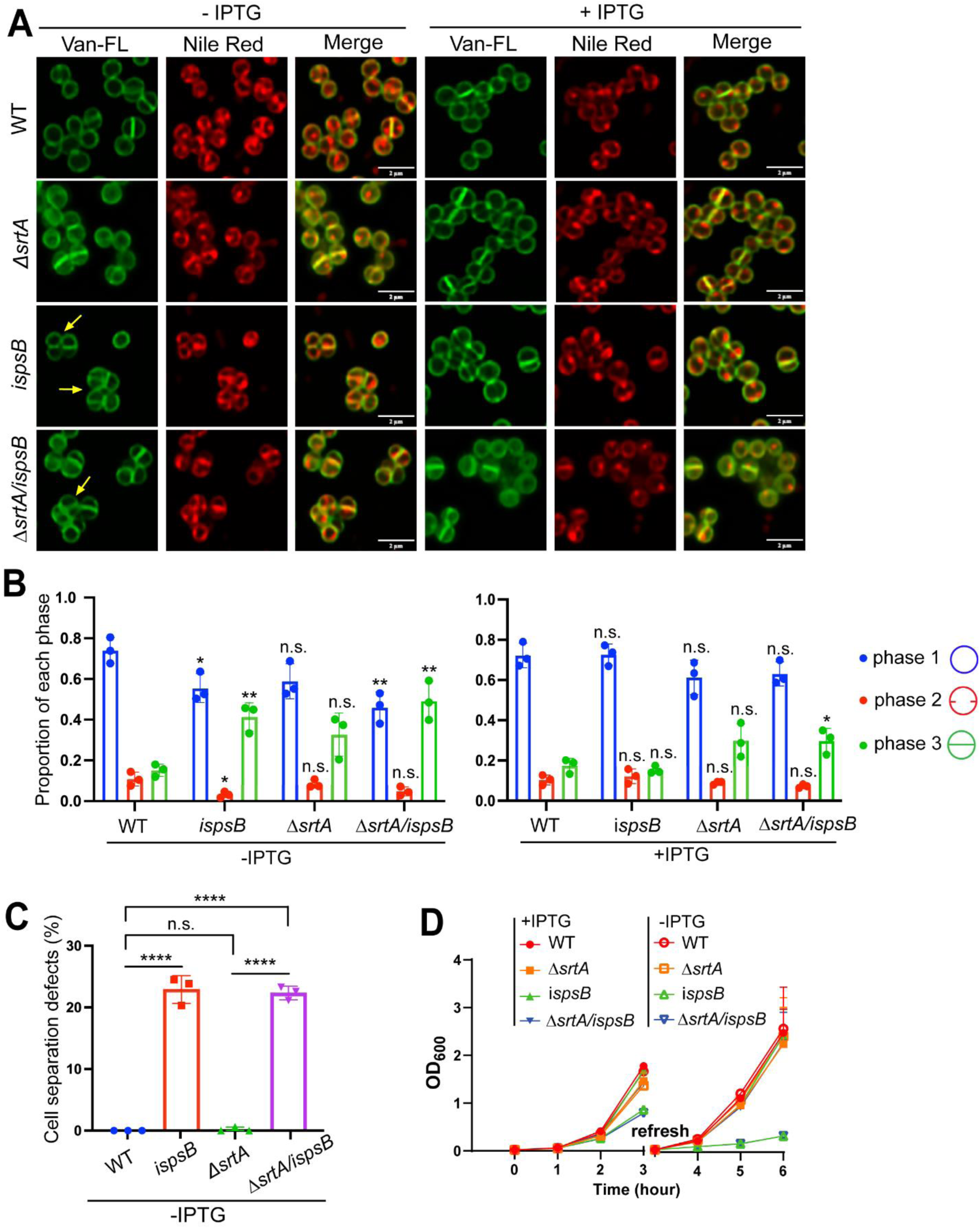
Depletion of *spsB* led to aberrant cell morphology, cell cycle arrest and growth defect. **(A)** Cell morphology of *spsB* depleted (-IPTG) and induced (+IPTG) cells. Fluorescent vancomycin (Van-FL) stains bacterial cell wall and Nile red stains cell membrane. Yellow arrows indicate cell septation and separation defects. Scale bar, 2 µm. **(B)** Quantification of cells from different stages of the cell cycle: with no septum (denoted as P1), a partial septum (denoted as P2), or a complete septum (denoted as P3). Asterisks on top of each sample indicate statistical analysis result between WT and the sample. **(C)** Quantification of cell separation defects based on Van-FL staining in panel A. Unpaired t-test with Welch’s correction was performed for statistical analysis in panel B and C: *p <0.05; **p <0.005; ***p <0.0005; ****p <0.0001. **(D)** Growth curves of *spsB* depleted (-IPTG) and induced (+IPTG) cells. Bacterial cultures were grown −/+ IPTG for 3 hours, refreshed and grown for another 3 hours.

The morphological defects of the *spsB* mutant promoted us to examine whether *spsB* depletion impaired cell wall biosynthesis. To address this, we performed a sequential fluorescent D-amino acids (FDAAs) incorporation experiment. FDAAs are actively incorporated into cell wall by penicillin-binding-proteins (PBPs) during cell division and growth, which have been used to monitor cell wall biosynthesis (Kuru *et al*, 2012; Kuru *et al*, 2019; Pereira *et al*, 2016). In our experiment, staphylococcal cells were incubated with HADA (blue), RADA (red) and OGDA (green) for 20 min sequentially and unbound FDAAs was washed away between rounds **(Fig. 5A)**. In the WT cells, HADA marked the oldest cell wall synthesis at the cell poles of two daughter cells, RADA marked the newer cell wall synthesis at the cross-wall between two daughter cells, and OGDA marked the newest cell wall synthesis at the new septa of two daughter cells (**Fig. 5A**). In comparison to WT and *srtA* mutant, a significant number (35-40%) of *spsB* depletion mutant cells showed delayed and irregular FDAAs incorporation, which appeared as punctuate foci at abnormal positions (indicated by yellow arrows in Fig. 5A and quantified in Fig. 5B, Fig. S3). These results indicate that cell wall biosynthesis is altered upon *spsB* depletion.

**Fig. 5.**
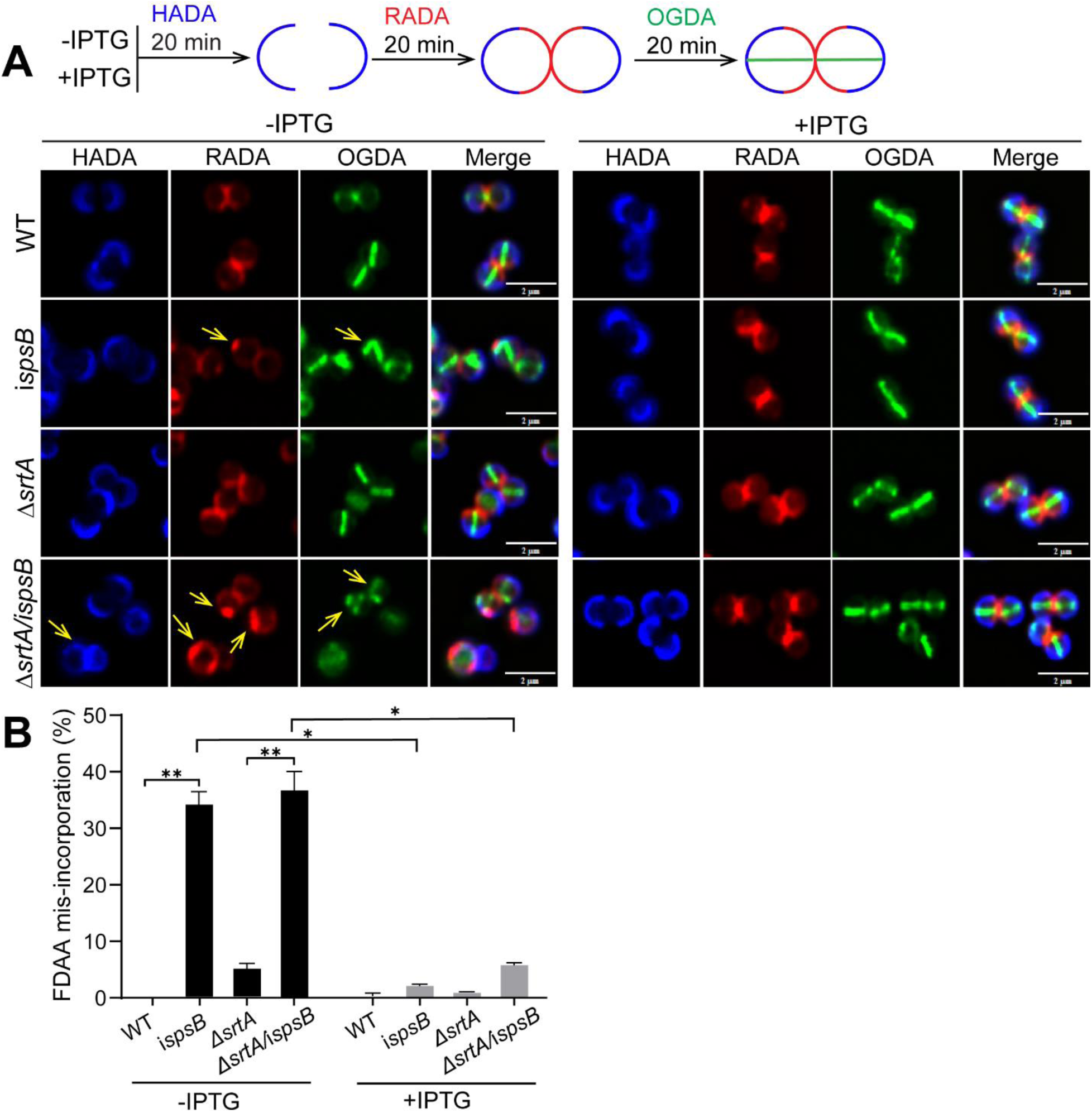
Depletion of *spsB* led to FDAA mis-incorporation. **(A)** *spsB*-depleted (-IPTG) and -induced (+IPTG) cells were sequentially incubated with HADA (blue), RADA (red) and OGDA (green). Yellow arrows indicate aberrant FDAA incorporation. Scale bar, 2 µm**. (B)** Quantification of FDAA mis-incorporation. Representative images and quantification are from three independent experiments. Unpaired t-test with Welch’s correction was performed for statistical analysis: *p <0.05; **p <0.005; ***p <0.0005; ****p <0.0001.

### SpsB predominantly localizes at the septum of dividing staphylococcal cells

Next, we sought to investigate the subcellular localization of SpsB in *S. aureus*. SpsB contains an N-terminal transmembrane domain and a C-terminal extracellular enzymatic domain (Madsen & Yu, 2024). Full-length SpsB was fused with N-terminal mCherry and expressed in the *spsB* depletion mutant. SpsB lacking its transmembrane domain (SpsB_Δ2-27_) fused with mCherry and SpsB alone were constructed as controls (**Fig. 6A**). Intriguingly, mCherry-SpsB was found primarily at the septum of dividing staphylococcal cells (**Fig. 6B**). In non-dividing cells, mCherry-SpsB was distributed all over the cell membrane. In comparison, mCherry-SpsB_Δ2-27_ failed to localize to the membrane but instead diffused in the cytoplasm. The localization of SpsB was quantified by calculating the fluorescence ratio (FR) of septal versus peripheral fluorescence signals (**Fig. 6C**). mCherry-SpsB signals were significantly increased at the septum compared to the non-specific membrane dye Nile red. Both mCherry-SpsB and mCherry-SpsB_Δ2-27_ produced intact fusion proteins as revealed by immunoblotting (**Fig. S4)**. However, expression of mCherry-SpsB or SpsB alone, but not mCherry-SpsB_Δ2-27_, rescued the growth defect, cell cycle retardation and cell separation defects caused by *spsB* depletion, indicating that mCherry-SpsB, but not mCherry-SpsB_Δ2-27_, was functional (**Fig. S4**). Taken together, we concluded that SpsB predominantly localized at the septum of dividing staphylococcal cells.

**Fig. 6.**
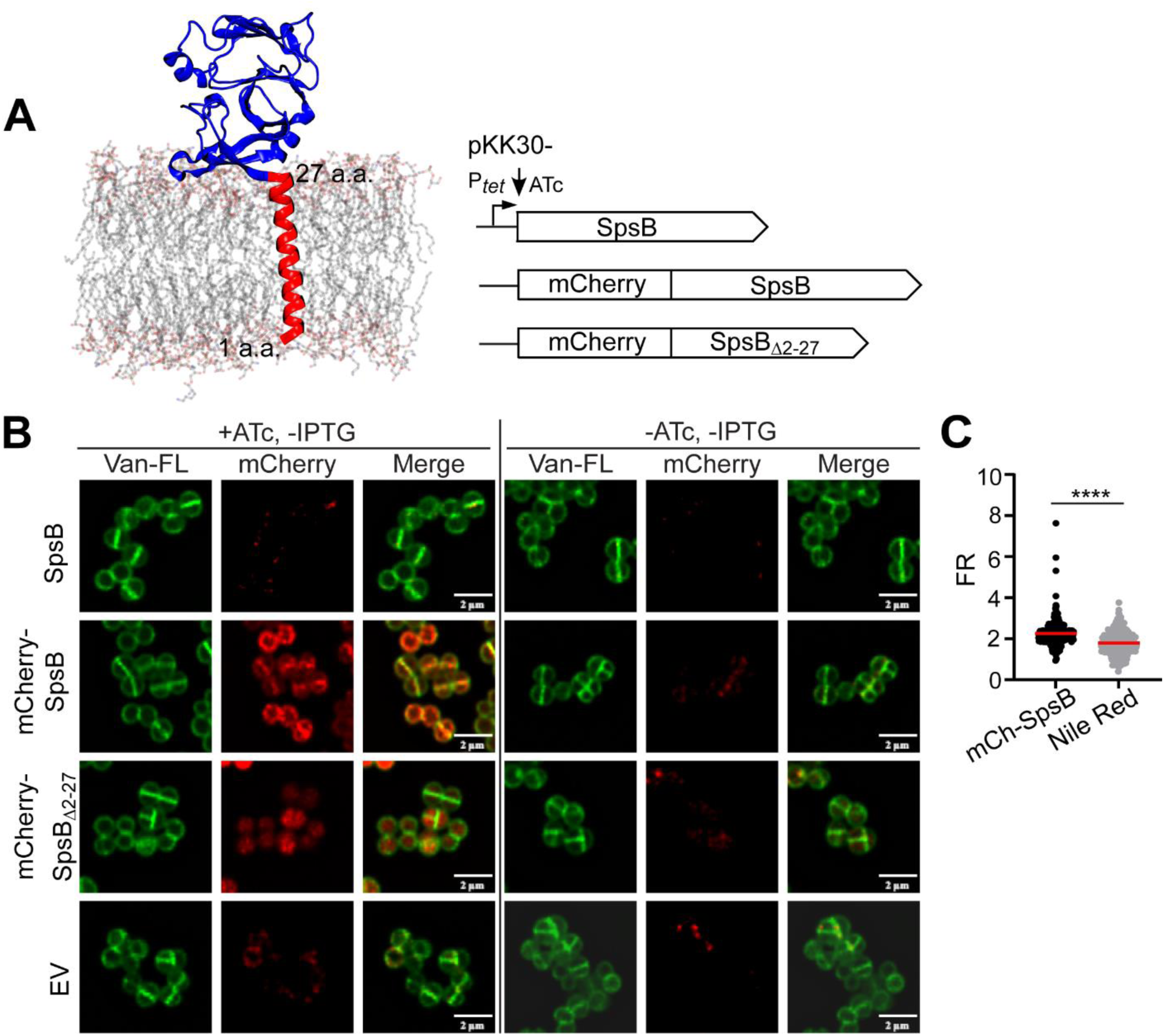
SpsB predominantly localizes at the septum of dividing staphylococcal cells. **(A)** Left: structural model of SpsB in the cell membrane generated by molecular dynamics simulation. Right: SpsB alone, mCherry fused with SpsB or SpsB lacking its transmembrane domain (SpsB_Δ2-27_) were expressed under ATc-inducible P*_tet_* promoter in SEJ1i*spsB*. **(B)** Fluorescence microscopy images showing the localization of mCherry-SpsB and mCherry-SpsB_Δ2-27_ in *spsB*-depleted cells (-IPTG). **(C)** Quantification of fluorescence intensity ratio (FR) of septum versus cell periphery. Unpaired t-test with Welch’s correction was performed for statistical analysis: *p <0.05; **p <0.005; ***p <0.0005; ****p <0.0001. Representative images and quantification are from three independent experiments.

### SpsB spatially regulates LtaS by cleaving LtaS at the septum

Our previous studies showed that LtaS-mediated LTA synthesis regulates SpA septal localization (Yu *et al*., 2018; Zhang *et al*., 2021). LtaS is cleaved by SpsB between Ala_217_ and Ser_218_, a position between its transmembrane domains and eLtaS (Wormann *et al*., 2011). LtaS has been shown to localize at the septum (Reichmann & Grundling, 2011). However, the localization of LtaS was revealed by GFP fusion to LtaS_S218P_, a point mutant of LtaS that is resistant to SpsB-mediated cleavage. GFP tagged wild-type LtaS was reported to be unstable (Reichmann & Grundling, 2011). Based on our results here that SpsB is enriched at the septum, we hypothesized that LtaS is more rapidly cleaved at the septum by SpsB; consequently, the LtaS_S218P_ variant accumulates at the septum. If this were true, we would expect that *spsB* depletion would enrich GFP-LtaS_WT_ fusion at the septum. To test it, we obtained the GFP-LtaS_S218P_ construct from the Gründling lab and generated GFP-LtaS_WT_ fusion to be expressed in the *spsB* depletion mutant. Consistent with the previous study (Reichmann *et al*., 2014), we were able to reproduce the result that GFP-LtaS_S218P_ localized at the septum in *ltaS*-depleted cells (**Fig. 7A**, GFP-LtaS_S218P_, i*ltaS*, -IPTG) and in SEJ1 WT (data not shown). Moreover, signals of GFP-LtaS_WT_ were diffused in the cytoplasm in *spsB*-induced cells (Fig. 7A, GFP-LtaS_WT_, i*spsB*, +IPTG), suggesting that the fusion protein was unstable. Finally, similar to the septal localization of GFP-LtaS_S218P_, depletion of *spsB* enriched GFP-LtaS_WT_ signal at the septum (Fig. 7A, GFP-LtaS_WT_, i*spsB*, -IPTG), although the signal was weaker than that of GFP-LtaS_S218P_.

**Fig. 7.**
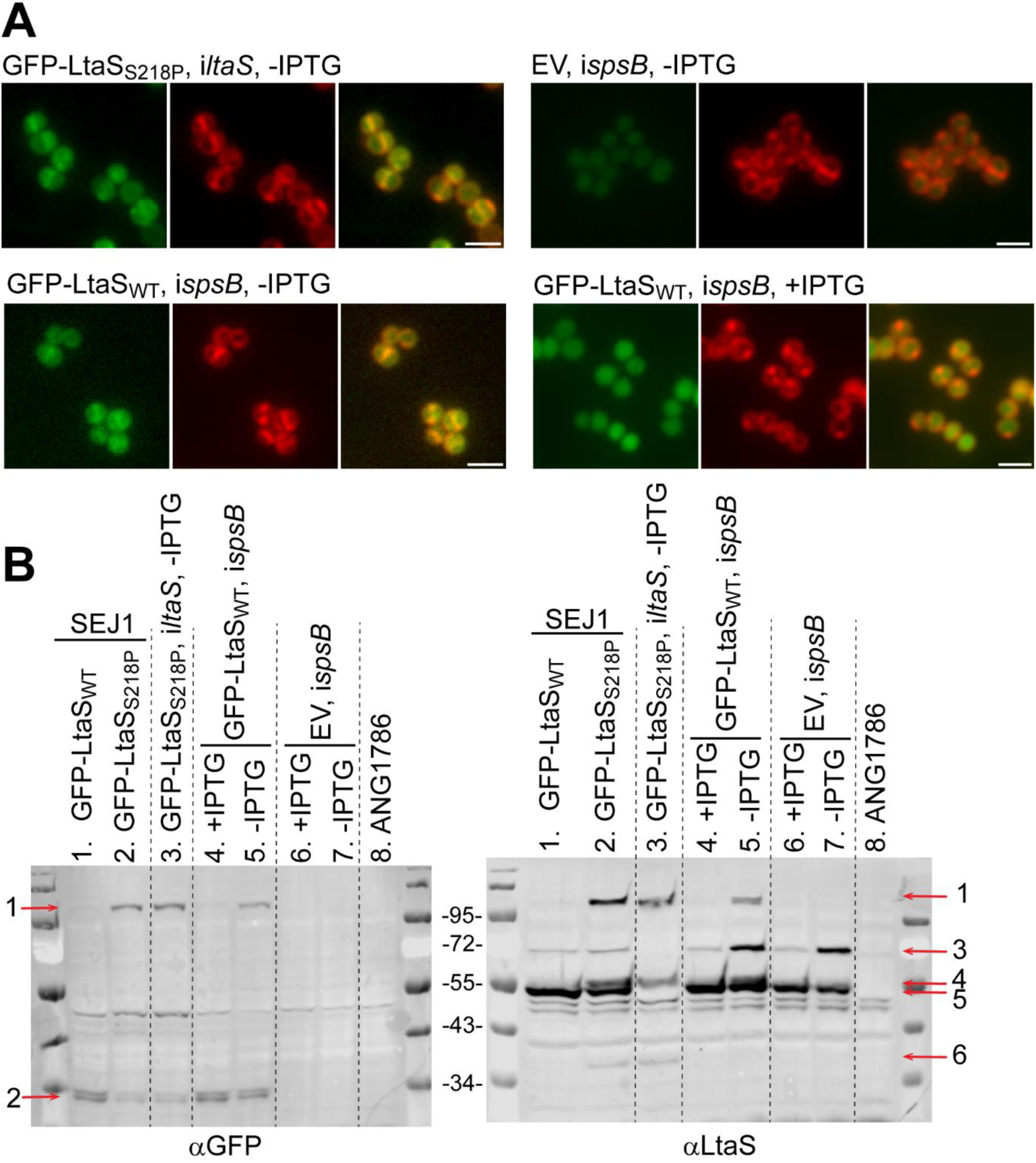
LtaS is enriched at the septum upon *spsB* depletion. (**A).** Fluorescence microscopy images showing GFP-LtaS_WT_ and GFP-LtaS_S218P_ localization in *ltaS* or *spsB*-depleted cells. Bacterial cultures were grown −/+IPTG and +ATc for 3 hours. EV: empty vector. Cell membrane was stained with Nile red. Scale bar: 2 µm. **(B)**. Anti-GFP and anti-LtaS immunoblot analysis of whole cell lysate. Sample order: Lane 1, GFP-LtaS_WT_ expressed in SEJ1 WT; lane 2, GFP-LtaS_S218P_ expressed in SEJ1 WT; lane 3, GFP-LtaS_S218P_ expressed in *ltaS*-depleted cells; lane 4, GFP-LtaS_WT_ expressed in *spsB*-induced cells; lane 5, GFP-LtaS_WT_ expressed in *spsB*-depleted cells; lane 6, empty vector control in *spsB*-induced cells; lane 7, empty vector control in *spsB*-depleted cells; lane 8, ANG1786, an *ltaS* deletion mutant as a negative control for GFP and LtaS blots. Red arrows indicate protein bands: arrow 1, full-length GFP-LtaS_WT/S218P_ fusion (theoretical MW: 102.5 kD); arrow 2, GFP-immunoreactive degradation products; arrow 3, native full-length LtaS (theoretical MW: 74.4 kD); arrow 4, GFP-LtaS_S218P_ degradation products in lane 2&3; arrow 5, presumably eLtaS (theoretical MW: 49.3 kD); arrow 6, GFP-LtaS_S218P_ degradation products in lane 2&3. Protein ladders in kDa are indicated between the blots.

We then performed anti-GFP and anti-LtaS immunoblotting to examine the expression and cleavage of GFP-LtaS_WT_, GFP-LtaS_S218P_ and native LtaS in our strains (**Fig. 7B**). An intact full-length fusion protein band was detected in SEJ1 WT or *ltaS*-depleted cells expressing GFP-LtaS_S218P_ (indicated by red arrow 1, lane 2&3 in both blots). The intact full-length fusion protein could also be detected in *spsB*-depleted cells expressing GFP-LtaS_WT_ (lane 5 in both blots), but the signals were weaker than that of lane 2 and 3. Concurrently more GFP-immunoreactive degradation products were found in lane 5 (indicated by red arrow 2 in αGFP blot). No full-length GFP-LtaS fusion bands could be detected in SEJ1 WT or *spsB* induced cells expressing GFP-LtaS_WT_ indicating rapid cleavage and degradation, which is in agreement with previous results (Wormann *et al*., 2011). Moreover, *spsB* depletion enriched full-length native LtaS (indicated by red arrow 3, lane 5&7 compared to lane 4&6 in αLtaS blot). The immunoblot results were consistent with the microscopy observations: GFP-LtaS_S218P_ produced more stable fusion protein that accumulated at the septum; depletion of *spsB* reduced cleavage and enriched fusion protein at the septum. Overall, the results support the hypothesis that SpsB preferentially cleaves LtaS at the septum, which potentially contributes to spatial regulation of LtaS-mediated LTA synthesis and consequently YSIRK+ protein septal trafficking.

## Discussion

In summary, the current study unveiled novel functions of the signal peptidase SpsB in regulating septal trafficking of YSIRK+ proteins, staphylococcal cell cycle and LtaS localization.

The first major finding is that the septal localization of SP_SpA(YSIRK+)_-SpA* is dependent on SpsB-mediated cleavage. SP_SpA_-SpA* lost its septal localization but dispersed all over the cell when *spsB* was depleted. SpsB is essential for bacterial viability and depletion of *spsB* led to aberrant cell morphology and defects in cell cycle, which could interfere with SpA* localization. However, the mis-localization was also observed with SP_SpA_A37P_-SpA* expressed in WT and Δ*srtA* with normal cell morphology. The results of SP_SpA_A37P_ demonstrated that the effect of SpsB is due to SP cleavage, but not related to aberrant cell morphology upon *spsB* depletion.

Consistent with previous proteomics data (Schallenberger *et al*, 2012), our experimental data showed that SpsB cleaves both SP_SpA(YSIRK+)_ and SP_SplE(YSIRK-)_. However, SpsB only affected the septal localization of SP_SpA(YSIRK+)_-SpA*, but not the peripheral localization of SP_SplE(YSIRK-)_-SpA*. A key question is, how would SpsB only affect the localization of SP_SpA(YSIRK+)_ but not SP_SplE(YSIRK-)_? We propose that 1) the YSIRK+ preproteins are more efficiently processed by SpsB at the septum leading to SpA* accumulation; 2) the cleavage efficiency is the same for both YSIRK+ and YSIRK-substrates, however YSIRK+ preproteins are more abundant and dose-wise more processed at the septum. These two possibilities are not necessarily mutually exclusive. Indeed, previous studies have shown that the YSIRK/G-S motif is required for efficient secretion and many YSIRK+ proteins are highly abundant (Bae & Schneewind, 2003; DeDent *et al*., 2008; Yu *et al*., 2018), which supports that both processing efficiency and preprotein abundance can play a role. We previously showed that SecA is the cytoplasmic transporter of SpA and SecA localized circumferentially along the cell membrane. Based on the results from this study, we propose a refined model whereby both YSIRK+ and YSIRK-preproteins bind to SecA and are transported all over the cell; the YSIRK+ preproteins are more abundant and/or more efficiently processed by SpsB at the septum, leading to the accumulation of YSIRK+ precursors at the septum. When the signal peptide cleavage is impaired by either *spsB* depletion or mutation in the SP cleavage site, YSIRK+ preproteins no longer accumulate at the septum but diffuse all over the membrane.

The second major finding is that SpsB predominantly localizes at the septum and regulates staphylococcal cell cycle. There are accumulating data from the literature showing that protein biogenesis and secretion apparatuses have specific cellular localizations that are linked to cell division and cell cycle. For example, SecA has been reported to localize at the septum of *Streptococcus pneumoniae* and *Streptococcus agalactiae* and as a single microdomain in *Streptococcus mutans* and *Enterococcus faecalis* (Brega *et al*, 2013; Hu *et al*, 2008; Kline *et al*, 2009; Tsui *et al*, 2011). The Sec translocon is organized in spiral-like structures in *Bacillus subtilis* (Campo *et al*, 2004). A recent study from *S. aureus* showed that the protein chaperone trigger factor is enriched at proximity to the septal membrane, which promotes the septal secretion of cell wall hydrolase Sle1 (Veiga *et al*, 2023). SecA is required for stalk biogenesis and cell division In *Caulobacter crescentus* (Kang & Shapiro, 1994). SecA drives transmembrane insertion of RodZ, the rod shape maintenance protein, and is essential for spatiotemporal organization of MreB, the bacterial actin homolog in *E. coli* (Govindarajan & Amster-Choder, 2017; Rawat *et al*, 2015). In *Bacillus subtilis*, SecA is required for membrane targeting of the cell division protein DivIVA (Halbedel *et al*, 2014). While most of the literature focused on SecA, little is known about SpsB. Interestingly, the localization and the mutant phenotypes of SpsB are distinctly different from that of SecA in *S. aureus:* SecA has a uniform localization*; secA-*depleted cells showed enlarged and irregular cell sizes but had no obvious cell separation or cell cycle defects (Yu *et al*., 2018). These results suggest a distinct role of SpsB in regulating staphylococcal cell cycle.

How does SpsB regulate staphylococcal cell cycle? Our data suggest that SpsB is a late-stage cell cycle regulator as the cell population at cell cycle phase 3 is significantly increased upon *spsB* depletion, suggesting that the cells are deficient in completing septum and daughter cell separation. Cell separation requires completion of cell wall synthesis and timely cleavage by cell wall hydrolases. The genome of *S. aureus* encodes more than a dozen cell wall hydrolases. LytN is the only cell wall hydrolases that contains a YSIRK+ signal peptide. It was shown previously that LytN is secreted at the septum and the mutant of *lytN* impaired cell separation (Frankel *et al*, 2011). Plausibly, *spsB* depletion alters the septal targeting of LytN, which leads to cell separation defects. Previous proteomics data showed that the secretion of autolysins Atl and Sle1 was reduced when *S. aureus* was treated with SpsB inhibitor arylomycin (Schallenberger *et al*., 2012). Although Atl and Sle1 are not YSIRK+ proteins, it is possible that SpsB processes Atl and Sle1 at the septum during cell division, which maximizes their activity; depletion of *spsB* decreases the secretion of Atl and Sle1 resulting in cell separation defects. Our experimental results from FDAA staining indicate that *spsB* depletion impaired cell wall biosynthesis. Whether it is the consequence of cell wall hydrolases dysregulation or direct impact from loss of SpsB activity remains to be addressed. The underlying mechanisms of how SpsB localizes at the septum and regulates cell cycle are currently under investigation.

The third major finding of this study is that SpsB spatially regulates LtaS, which potentially regulates LTA synthesis and modulates septal trafficking of YSIRK+ proteins. LtaS is an unusual substrate of SpsB. The cleavage of SpsB separates eLtaS from its transmembrane domains. The functional relevance of LtaS cleavage by SpsB remains unclear. It was previously reported that the cleavage of LtaS by SpsB inactivates its enzymatic activity (Wormann *et al*., 2011). A recent study observed LtaS cleavage and LTA production over time and found that the LtaS_S218P_ mutant was delayed in eLtaS release and LTA production. It was proposed that SpsB-mediated cleavage temporally regulates LtaS activity (Ibrahim *et al*., 2024). Another study using transposon mutagenesis with outward promoters revealed that reduced expression of LtaS confers high resistance to the SpsB inhibitor M131 (Meredith *et al*, 2012). The mechanism by which LtaS expression modulates resistance to SpsB inhibitor is unclear. Our results here suggest that the cleavage event is linked to the spatial regulation of LtaS. Interestingly, we showed previously that LTA is more abundant at the cell periphery than at the septum. We reasoned that mature LTA accumulates at the cell periphery over continuous rounds of cell division. Based on the results here, it is possible that LtaS is less active at the septum thereby generating less LTA at the septum. How does SpsB-mediated LtaS cleavage regulate septal trafficking of YSIRK+ proteins? A recent study showed that the LtaS_S218P_ mutant slowly releases eLtaS to the supernatant, which correlates with SpA cross-wall localization. Based on our results here, we propose that eLtaS is released at the septum, which spatially modulates SpA septal trafficking.

Taken together, we propose a new dual-mechanism model mediated by SpsB in regulating YSIRK+ surface protein trafficking (**Fig. 8**). In the WT cells, the highly expressed YSIRK+ proteins are more efficiently processed by SpsB at the septum, which enriches YSIRK+ protein precursors to be anchored to septal peptidoglycan. Moreover, SpsB cleaves LtaS and releases eLtaS at the septum, which further supports the septal trafficking of YSIRK+ proteins. In *spsB*-depleted cells, the cleavage of SP_YSIRK+_ and LtaS is impaired, resulting in dispersed localization of YSIRK+ proteins and accumulation of full-length LtaS at the septum. In addition, SpsB plays a key role in regulating staphylococcal cell cycle. Our study sheds light on novel molecular mechanisms coordinating protein secretion, cell cycle and cell envelope biogenesis in *S. aureus.* As SpsB, YSIRK/G-S signal peptides and LTA synthesis are widely distributed and conserved in Gram-positive bacteria, the mechanisms identified here may be applicable in other bacteria.

**Fig. 8.**
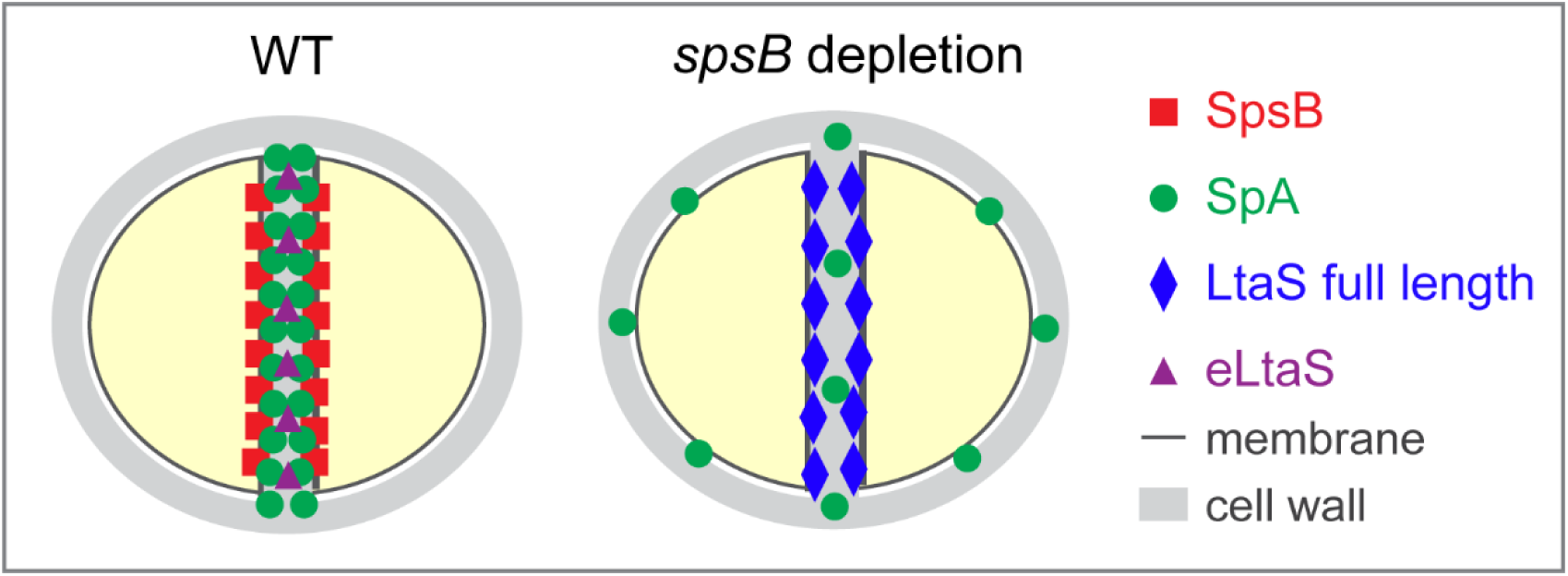
Model of how SpsB regulates the processing and localization of SpA (YSIRK+ protein) and LtaS. In wild-type *S. aureus* cells: SpsB efficiently cleaves SpA preproteins and full-length LtaS at the septum, leading to SpA accumulation and eLtaS release at the septum. In *spsB*-depleted cells: the cleavage of signal peptide and LtaS is impaired; SpA preproteins disperse all over the cell membrane and full-length LtaS accumulate at the septum.

## Materials and Methods

### Bacterial strains and growth conditions

Strains and plasmids used in this study are listed in Table S1. *Escherichia coli* strains were grown in Luria-Bertani broth (LB) or on LB agar plates. *S. aureus* strains were grown in tryptic soy broth (TSB) or on tryptic soy agar plates (TSA). For plasmid selection in *E. coli*, 100 µg/ml ampicillin (Amp) or 10 µg/ml trimethoprim (Tmp) was used. For plasmid and mutant selection in *S. aureus*, 5-10 µg/ml chloramphenicol (Chl), 10 µg/ml erythromycin (Ery), and 50 µg/ml kanamycin (Kan) were used. 1 mM isopropyl β-D-1-thiogalactopyranoside (IPTG) was used to induce *spsB* expression from the P*_spac_* promoter and 200 ng/ml anhydrotetracycline (ATc) was used to induce gene expression from the P*_tet_* promoter. If not specified, overnight cultures were grown with appropriate antibiotics and IPTG when needed. Overnight cultures were 1:100 diluted in fresh TSB with or without IPTG or ATc. Refreshed cultures were grown for 2-3 hours to mid-log phase (OD_600_ of 1.0) or to a desired OD_600_ for different experiments.

### Construction of strains and plasmids

To construct SEJ1*ΔsrtA*, the pKOR1 knock-out plasmid was used (Bae & Schneewind, 2006). Primers 673/674 and 675/676 were used to amplify the upstream and downstream fragments surrounding *srtA* of SEJ1. The PCR products were ligated via overlapping PCR and cloned into pKOR1 via BP clonase reaction to generate pKOR1-*srtA*. The plasmid was electroporated to SEJ1 and subjected to the integration process by incubation at 42°C for two rounds. Subsequently, cultures were streaked on TSA (Chl-5) plates and incubated at 42°C overnight. Several colonies were selected and grown in antibiotic-free TSB at 30°C overnight. Next, cultures were plated on TSA containing 200 ng/ml ATc and incubated at 37°C overnight. To exclude cointegrate contamination, clones were streaked on TSA with and without Chl-5; those that did not grow on TSA (Chl-5) were selected. Positive clones were verified by PCR with primers 677/678 and 679/680 and confirmed by DNA sequencing. SEJ1*ΔsrtAispsB* was constructed by first transforming SEJ1*ΔsrtA* with pCL55-P*_spac_-spsB*-T114C, followed by transducing *ΔspsB::erm*. SEJ1*srtA*::φΝΣ transposon mutant was generated by phage transduction from Nebraska Transposon Mutant Library (NTML).

To construct pCL*itet-sp_splE_-spa*,* primers 562/563 were used to amplify the signal peptide of *splE*. Mutagenesis PCR was used to replace *sp_spa_* with *sp_splE_* in plasmid pCL*itet-sp_spa_-spa\** (Zhang *et al*., 2021). The PCR products were digested with DpnI and transformed to *E. coli* DC10B (Monk *et al*, 2012). Similarly, site-directed mutagenesis PCR was used to construct pCL*itet-sp_spa_A37P_-spa\** using primers 416/417. To construct pKK30*itet-sp_spa_-spa\** and pKK30*itet-sp_splE_-spa\**, primers 681/682 were used to amplify P*itet-sp_spa_-spa\** and P*itet-sp_splE_-spa\** from template pCL*itet-sp_spa_-spa\** and pCL*itet-sp_splE_-spa\**. The PCR products and the vector pKK30 (Krute *et al*, 2016) were digested with NotI/SacI and ligated.

To construct pKK30*itet-spsB,* primers 972/973 were employed to amplify *spsB* from RN4220 genomic DNA and the vector pKK30*itet* was digested with XmaJI/BglII followed by Gibson Assembly (NEB) to generate pKK30*itet-spsB.* To construct mCherry-SpsB fusion plasmids, primers 948/887 were used to amplify *spsB* from RN4220 genomic DNA and primers 886/837 were used to amplify *mCherry* from pCX-gpmCh-cw1 (Yu & Götz, 2012). The vector pCL*itet* was digested by SalI/BglII followed by Gibson Assembly combining *spsB* and *mCherry* to generate pCL*itet-mCherry-spsB.* Next, primers 925/926 were used to amplify *mCherry-spsB* from pCL*itet-mCherry-spsB* and ligated with XbaI/SacI-digested pKK30*itet-sp_spa_-spa\** via Gibson Assembly to generate pKK30*itet-mcherry-spsB.* To construct pKK30*itet-mcherry-spsB_Δ2-27_,* primers 973/974 were employed to amplify *spsB_Δ2-27_* from RN4220 genomic DNA. The plasmid pKK30*itet-mcherry-spsB* was digested with NheI/BglII and ligated with the PCR product of *spsB_Δ2-27_* by Gibson Assembly.

To construct pKK30*itet*-*gfp_P7_*-*ltaS_WT_* and pKK30*itet*-*gfp_P7_*-*ltaS_S218P_*, we first generated pCL*itet-gfp_P7_-ltaS_WT_* from template pCL*itet-gfp_P7_-ltaS_S218P_* (Reichmann *et al*., 2014) by site-directed mutagenesis PCR using primers 1023/1024. Both pCL*itet-gfp_P7_-ltaS_S218P_* and pCL*itet-gfp_P7_-ltaS_WT_* were digested with XmaJI/BglII and ligated with plasmid pKK30*itet*-*mCherry*-*spsB* that were cleaved with the same enzymes. All the strains and plasmids constructed were confirmed by DNA sequencing. Primers used in this study are listed in Table S2.

### SpA immunofluorescence microscopy (IF) and quantification

Two protocols described previously were used: cross-wall IF and membrane IF (Scaffidi *et al*., 2021; Scaffidi & Yu, 2024). (1) Cross-wall IF was used to detect newly synthesized protein on the bacterial cell surface. 2 ml of mid-log phase cultures were harvested, washed once with PBS, and incubated in 1 ml PBS containing 0.5 mg/ml trypsin at 37°C with rotation for 1 hour to remove the pre-deposited proteins on the cell surface. Trypsinized cells were washed twice with PBS and grown in fresh TSB supplemented with 1 mg/ml of soybean trypsin inhibitor (Sigma) for 20 min to allow protein regeneration. Cells were immediately fixed, washed with PBS, resuspended in 150 µl of PBS and applied to poly-L-lysine coated slides. (2). Membrane IF was used to detect protein localization underneath cell wall. Trypsinized cells were washed, fixed and resuspended in 1 ml GTE buffer (50 mM glucose, 20 mM Tris-HCl pH 7.5, 10 mM EDTA). After adding 20 µg/ml lysostaphin (AMBI), 50 µl of cell suspension was immediately transferred to poly-L-lysine coated slides and incubated for 2 min. Non-adherent cells were sucked away by vacuum and the slides were air-dried. Dried slides were immediately dipped in methanol at −20°C for 5 min and acetone for 30 s. After the slides were completely dried, samples were rehydrated with PBS and underwent immunofluorescence microscopy.

For immunofluorescence microscopy: slides prepared above were blocked with 3% BSA and incubated with rabbit anti-SpA_KKAA_ antiserum (Kim *et al*, 2010) (1:4,000 in 3% BSA) overnight at 4°C. Cells were washed with PBS eight times and incubated with Alexa Fluor 488 conjugated anti-rabbit IgG (1:500 in 3% BSA) (Invitrogen) for 1 hour in the dark. Unbound secondary antibodies were washed away ten times with PBS and cells were incubated with 50 µg/ml Hoechst 33342 DNA dye, 5 µg/ml Nile red (Sigma) for 5 min in the dark, followed by washing with PBS. A drop of SlowFade™ Diamond Antifade Mountant (Invitrogen) was applied to the slide before sealing with the coverslip. Fluorescent images were captured on a Nikon Scanning Confocal Microscope Eclipse Ti2-E with HC PL APO 63×oil objective (1.4 NA, WD 0.14 mm). For image quantification: at least 300 cells for each sample from three independent experiments were analyzed by ImageJ (Schneider *et al*, 2012). The total cell numbers and cells displaying cross-wall/septal/peripheral SpA* localization or no signaling were counted. Unpaired t-test with Welch’s correction was performed for statistical analysis using GraphPad Prism; p-values < 0.05 were regarded as significant.

### Cell fractionation and immunoblotting

The protocols have been used previously (Yu *et al*., 2018; Zhang *et al*., 2021). To separate bacterial culture supernatant and cell pellet, 1 ml of mid-log phase culture was normalized to OD_600_ of 1.0 and then collected by centrifugation at 13,000 rpm for 5 min. The supernatant was transferred to a new tube. The pellet was resuspended in 1 ml of membrane buffer [50 mM Tris-HCl (pH 7.5), 150 mM NaCl] containing 20 µg/ml lysostaphin and incubated for 30 min at 37°C. Proteins from the supernatant and the cell lysate were precipitated with 10% TCA on ice for 30 minutes, washed with acetone, air-dried, and solubilized in 100 µl SDS sample buffer. [62.5 mM Tris-HCl (pH 6.8), 2% SDS, 10% glycerol, 5% 2-mercaptoethanol, 0.01% bromophenol blue]. To separate cytosolic (C), membrane (M), cell wall (W), supernatant (S) fractions, bacterial cultures were normalized to OD_600_ of 1.0 in 1 ml TSB and centrifuged at 13,000 rpm for 5 min. The culture supernatant was transferred to Tube 1. The pellet was resuspended in 1 ml of TSM buffer [50 mM Tris-HCl (pH 7.5), 0.5 M sucrose, 10 mM MgCl_2_] containing 20 µg/ml lysostaphin and incubated at 37°C for 10 min. Subsequently, the cell lysate was centrifuged at 14,000 rpm for 5 min and the supernatant (cell wall fraction) was transferred to a new tube (Tube 2). The pellet was washed with 1 ml of TSM and resuspended in 1 ml of membrane buffer, followed by five times of freeze-thaw cycles in dry ice/ethanol and a warm water bath. Membrane fractions were sedimented by centrifugation at 14,000 rpm for 30 min, and the supernatant (cytosolic fraction) was transferred to Tube 3, while the pellet (membrane fraction) was resuspended in 1 ml of membrane buffer (Tube 4). Proteins from Tubes 1-4 were precipitated by 10% TCA, washed with acetone, air-dried and solubilized in 100 µl of SDS sample buffer. To collect proteins from the whole cell culture, bacterial cultures were normalized to OD_600_ of 1.0 in 1 ml and incubated with 20 µg/ml lysostaphin at 37°C for 30 min. The whole culture lysate was precipitated with 10% TCA, washed with acetone, air-dried and solubilized in 100 µl of SDS sample buffer. For immunoblotting, protein samples were separated by 10% or 12% SDS-PAGE and transferred onto polyvinylidene difluoride membranes. Membranes were blocked with 5% nonfat milk (supplemented with 0.25% human IgG to block SpA if needed) and probed with primary antibodies (αSpA_KKAA_ 1:20,000, αSrtA 1:20,000, αGFP 1:5,000, αmCherry 1:1,000, αLtaS 1:5,000), followed by incubation with secondary antibodies of IRDye 680LT goat anti-rabbit at 1:20,000 dilution. Immunoblots were scanned by Li-Cor Odyssey CLx 9140.

### Fluorescence microscopy

Depletion of *spsB*: all strains were incubated overnight in TSB supplemented with IPTG. Next morning, bacteria cultures were washed twice with fresh TSB and inoculated at OD_600_ of 0.05 with and without IPTG. Bacteria were grown for 3 hours, diluted again in fresh TSB to OD_600_ of 0.05 and grown for another 3 hours (6 hours of depletion). Samples were taken to measure OD_600_ and subjected to microscopy experiments at different time points. To visualize the cell morphology of the *spsB* depletion mutant, mCherry-SpsB and GFP-LtaS localization, bacterial cultures at desired growth phase were harvested, washed once with PBS, resuspended in 250 µl of PBS, mixed with fixation solution (2.5% paraformaldehyde and 0.01% glutaraldehyde in PBS) and incubated at room temperature for 20 min. Fixed cells were washed with PBS twice, resuspended in 150 µl of PBS and applied to poly-L-lysine coated slides. Samples were stained with 50 µg/ml Hoechst 33342 DNA dye, 5 µg/ml Nile red (Sigma), and 1 µg/ml fluorescent Vancomycin (Vancomycin-BODIPY, Van-FL) for 5 min followed by 3 times wash with PBS. A droplet of SlowFade™ Diamond Antifade Mountant was applied to each sample prior to sealing with coverslips. Fluorescent images were acquired using the Nikon Scanning Confocal Microscope ECLIPSE Ti2-E with HC PL APO 63×oil objective.

Quantification of cell cycle and cell separation defects and statistical analysis: at least 300 cells for each sample from three times independent experiments were analyzed in ImageJ. Van-FL images were used to quantify the cell cycles. The staphylococcal cell cycle has been defined previously (Monteiro *et al*, 2015): cells in phase 1 with no septum; phase 2, partial septum; phase 3, complete septum and elongated shape. To quantify cell separation defects, Van-FL images were used and the proportion of cells with a septation defect were calculated. Unpaired t-test with Welch’s correction was performed for statistical analysis using GraphPad Prism.

Quantification and statistical analysis of fluorescence intensity ratio (FR): The method has been described previously (Atilano *et al*, 2010; Yu & Götz, 2012; Zhang *et al*., 2021). Images from three independent experiments containing at least 300 cells with complete septa were included for quantification. To quantify the fluorescence ratio of septum versus periphery, a line was drawn perpendicular to and across the middle of the septum. Signal intensities at the septum and cell poles were measured in ImageJ. For comparison, the same bacterial samples were stained with Nile red, and the FR was quantified. Unpaired t-test with Welch’s correction was performed for statistical analysis using GraphPad Prism.

### FDAA incorporation and quantification

Bacterial cultures (with and without IPTG) were normalized to OD_600_ of 1.0, resuspended in 150 µl of TSB containing 1 mM HADA (Tocris Bioscience) and incubated at 37°C for 20 min. The samples were washed once with 500 µl of PBS, resuspended in 150 µl of TSB containing 0.5 mM RADA and incubated at 37°C for 20 min. The same procedure was repeated once more with 1 mM OGDA in TSB. Samples were washed twice with 500 µl of PBS, resuspended in 150 µl of PBS and loaded to poly-L-lysine coated slides. Fluorescent images were acquired using Keyence microscope BZ-X710 or Nikon Scanning Confocal Microscope Eclipse Ti2-E. Abnormal FDAA localizations were quantified in ImageJ. Statistical analysis of unpaired t-test with Welch’s correction was performed using GraphPad Prism.

### Molecular dynamics simulation

Structural model of SpsB in the Gram-positive bacteria phospholipid membrane. This model was generated by combining the crystal structure of the apo form of the peptidase (PDB ID: 4WVG) (Ting *et al*, 2016) and the transmembrane segment from the AlphaFold2-predicted structure corresponding to UniProtKB entry A0A0H3KFC9 (Jumper *et al*, 2021; Varadi *et al*, 2022). SpsB was embedded in the Gram-positive bacteria membrane [60:35:5 ratio of PG, lysyl-PG, and cardiolipin, a widely used composition (Kiirikki *et al*, 2024; Mohanan *et al*, 2020)] and subjected to minimization and a short molecular dynamics simulation. For simulation details, refer to (Madsen & Yu, 2024).

## Acknowledgements

This work is supported by NIH/NIGMS-R35GM146993 and the start-up funds from University of South Florida to WY. We thank Angelika Gründling for generously providing strains ANG2009 and GFP-LtaS_S218P_. We thank members of the Yu lab for critically reading the manuscript and providing suggestions.

---

**Fig. S1.**
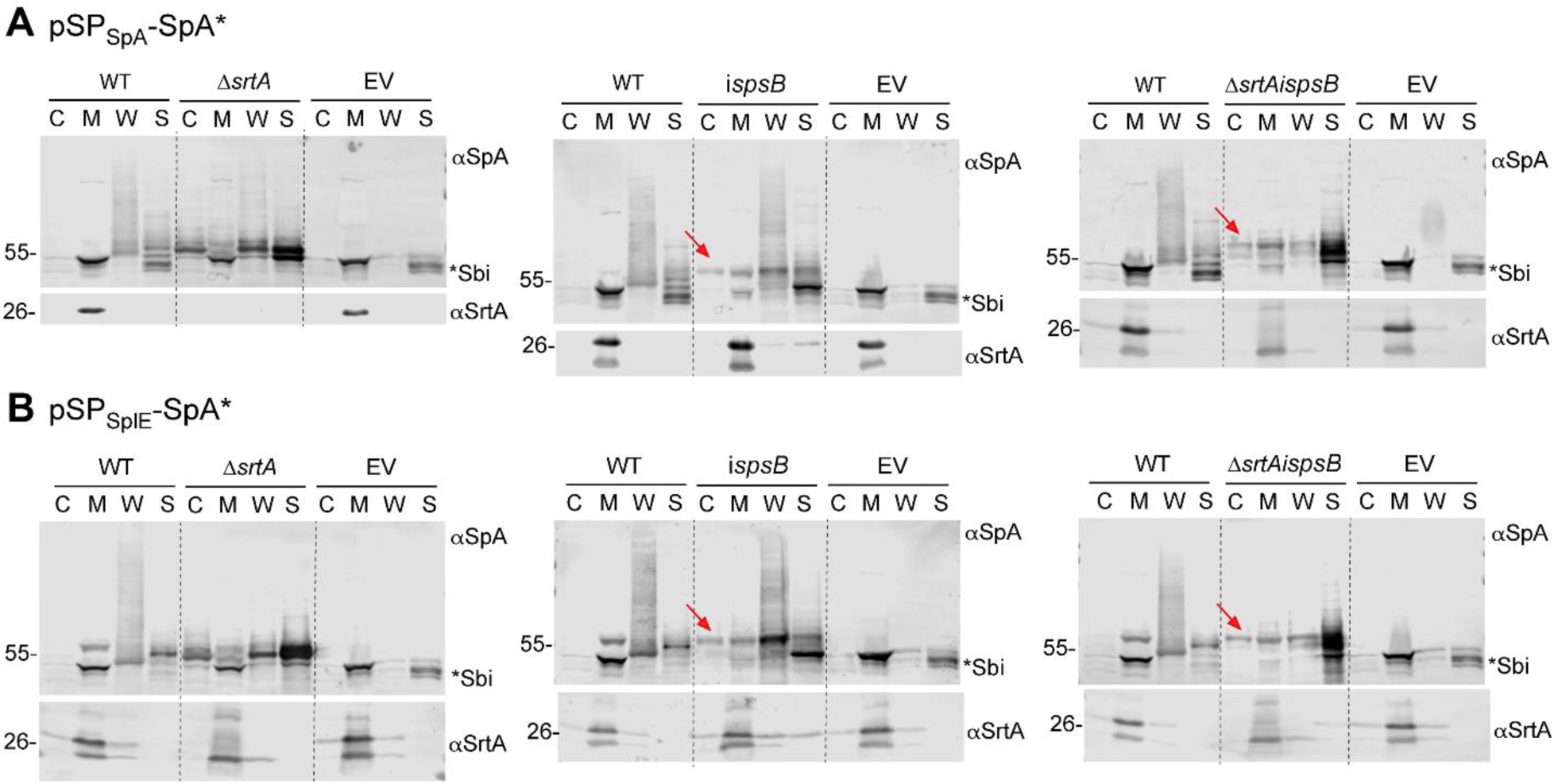
Cell fractionation and immunoblot analysis SpA* with empty vector (EV) control. Bacterial cultures of SEJ1 WT, Δ*srtA*, *ispsB*, Δ*srtA/ispsB* expressing SpA* fused with SP_SpA_ or SP_SplE_ were fractionated to cytoplasm (C), cell membrane (M), cell wall (W), and the supernatant (S). All the strains were grown without IPTG to deplete *spsB* and with ATc to induce *spa*\* expression. The αSrtA blot is a loading and fractionation control. The red arrow indicates unprocessed SP-bearing precursors. The asterisk indicates non-specific Sbi bands. Numbers on the left indicate protein ladder in kDa.

**Fig. S2.**
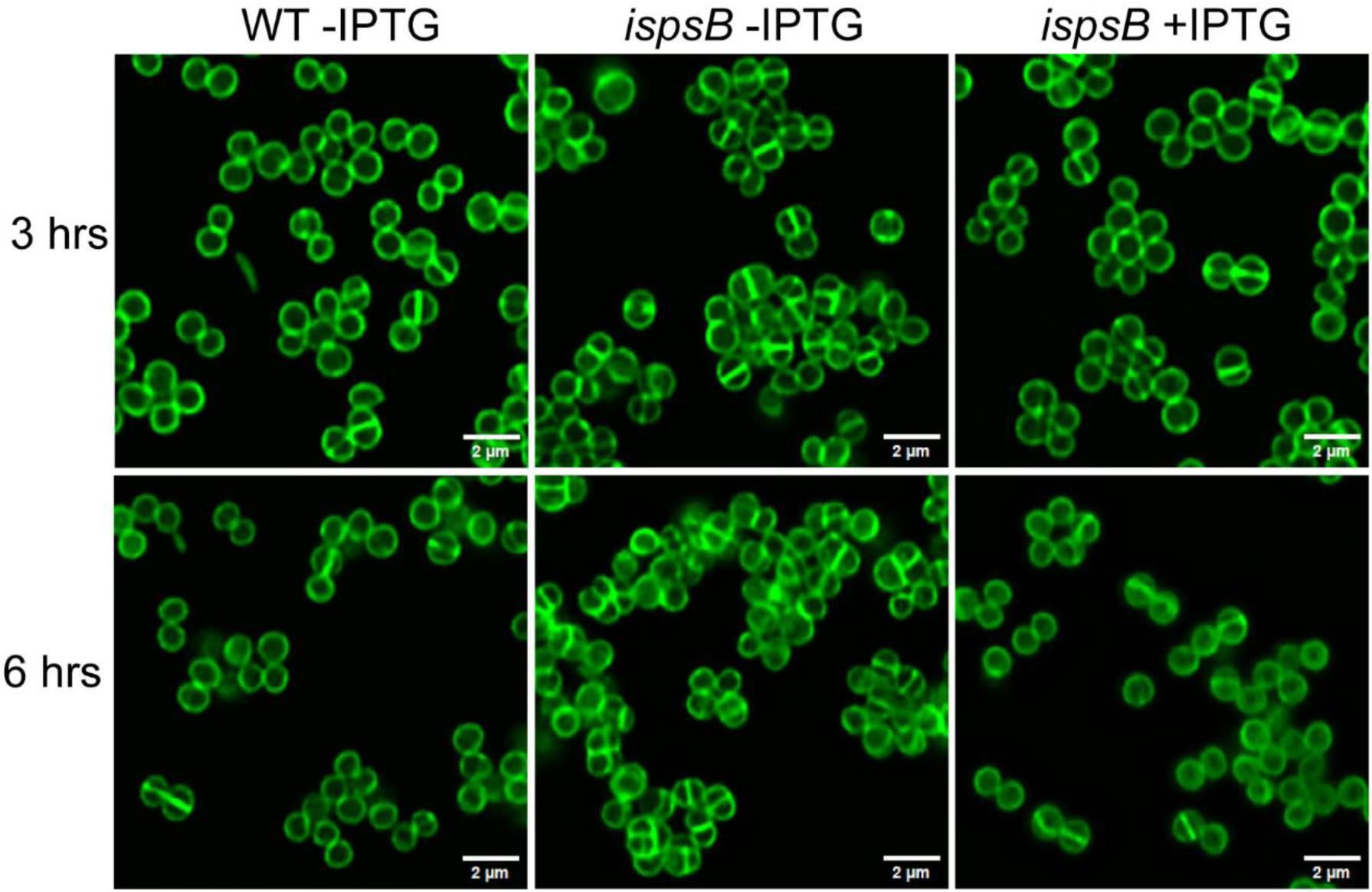
Extended *spsB* depletion led to elevated cell cycle arrest. Staphylococcal cells were stained with Van-FL after 3- and 6-hours of *spsB* depletion.

**Fig. S3.**
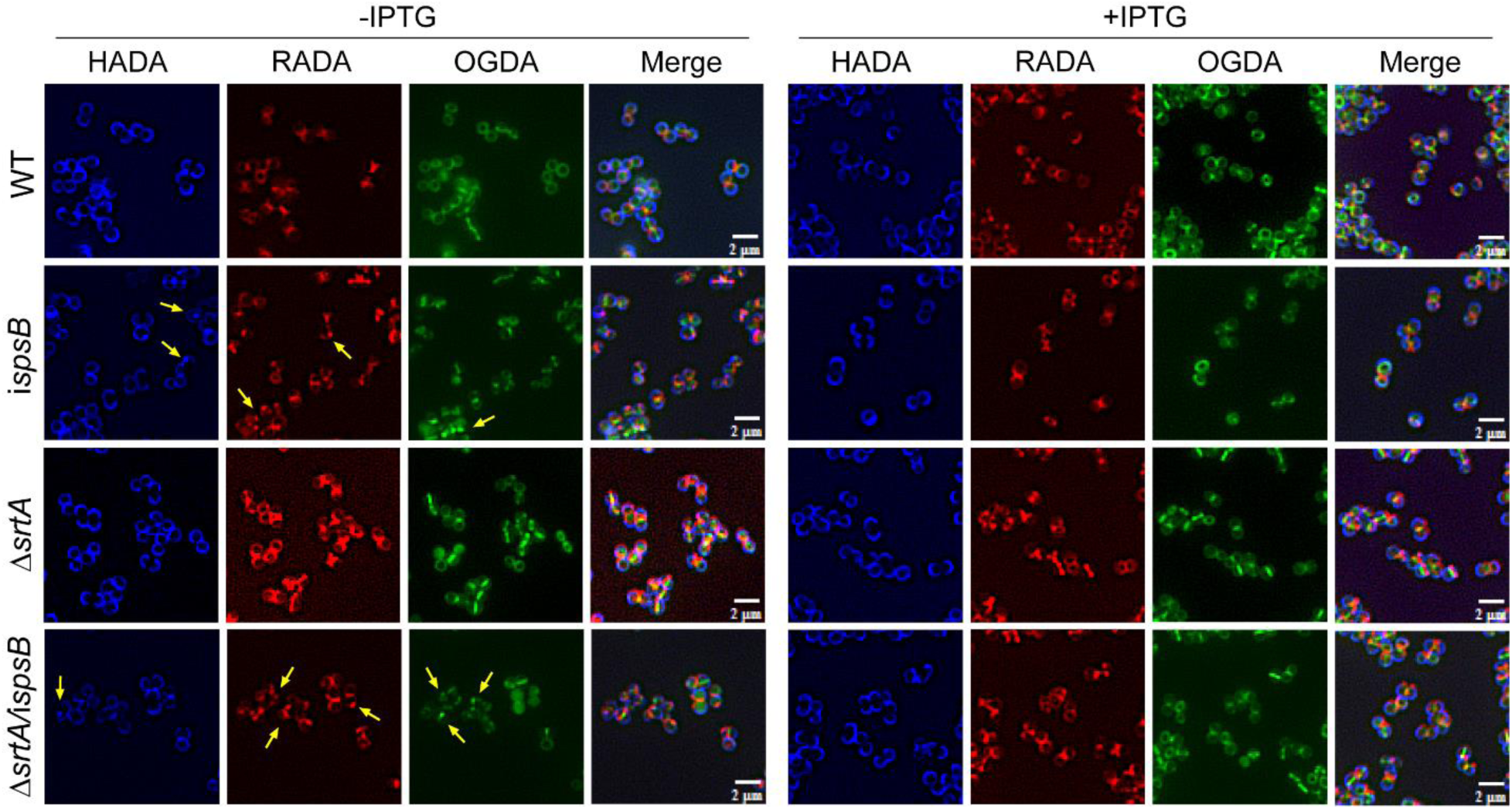
Extended data figure of Fig. 5. Larger image crops showing defects of FDAA incorporation upon *spsB* depletion, indicated by yellow arrows.

**Fig. S4.**
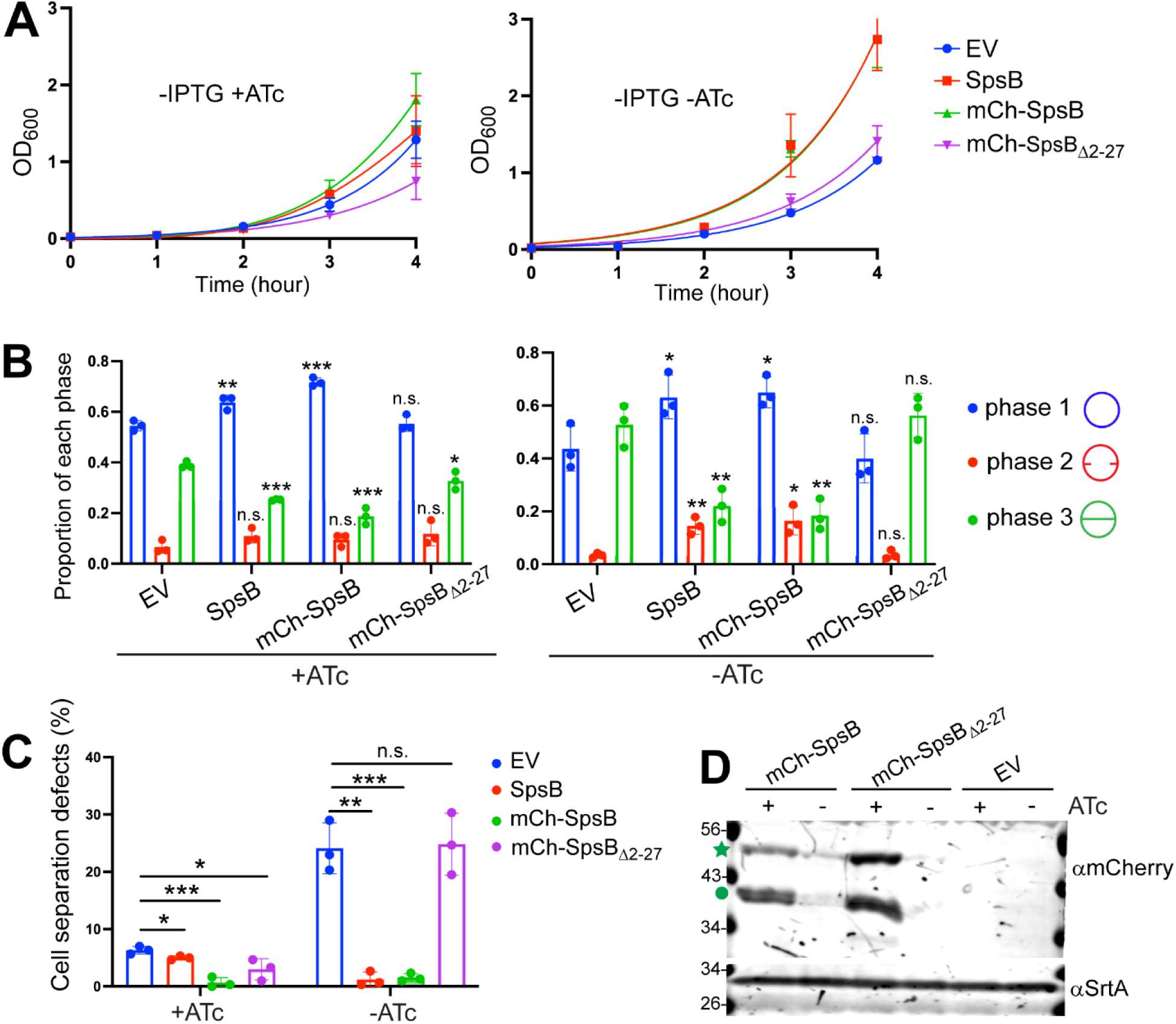
The mCherry-SpsB fusion is functional. **(A)** Growth curves of SEJ1*ispsB* expressing pKK30*itet* empty vector (EV), SpsB, mCherry-SpsB, mCherry-SpsB_Δ2-27_. Bacterial cultures were grown without IPTG to deplete *spsB* and with or without ATc to control the expression of the fusion proteins. **(B)** Quantification of cells from different stages of the cell cycle: with no septum (denoted as P1), a partial septum (denoted as P2), or a complete septum (denoted as P3). Asterisks on top of each sample indicate statistical analysis result between EV and the sample: *p <0.05; **p <0.005, ***p <0.0005; ****p <0.0001. **(C)** Quantification of cell separation defect based on Van-FL staining in Fig.6B. Unpaired t-test with Welch’s correction was performed for statistical analysis in panel B and C: *p <0.05; **p <0.005; ***p <0.0005; ****p <0.0001. **(D)** Anti-mCherry immunoblot analysis of whole cell culture. mCherry-SpsB, theoretical MW: 48.8 kD; mCherry-SpsB_Δ2-27_, theoretical MW: 45.8 kD. The star indicates intact fusion protein. Circle indicates degradation products. The αSrtA blot serves as a loading control. The protein ladders in kDa are noted on the left side.

**Table S1.**
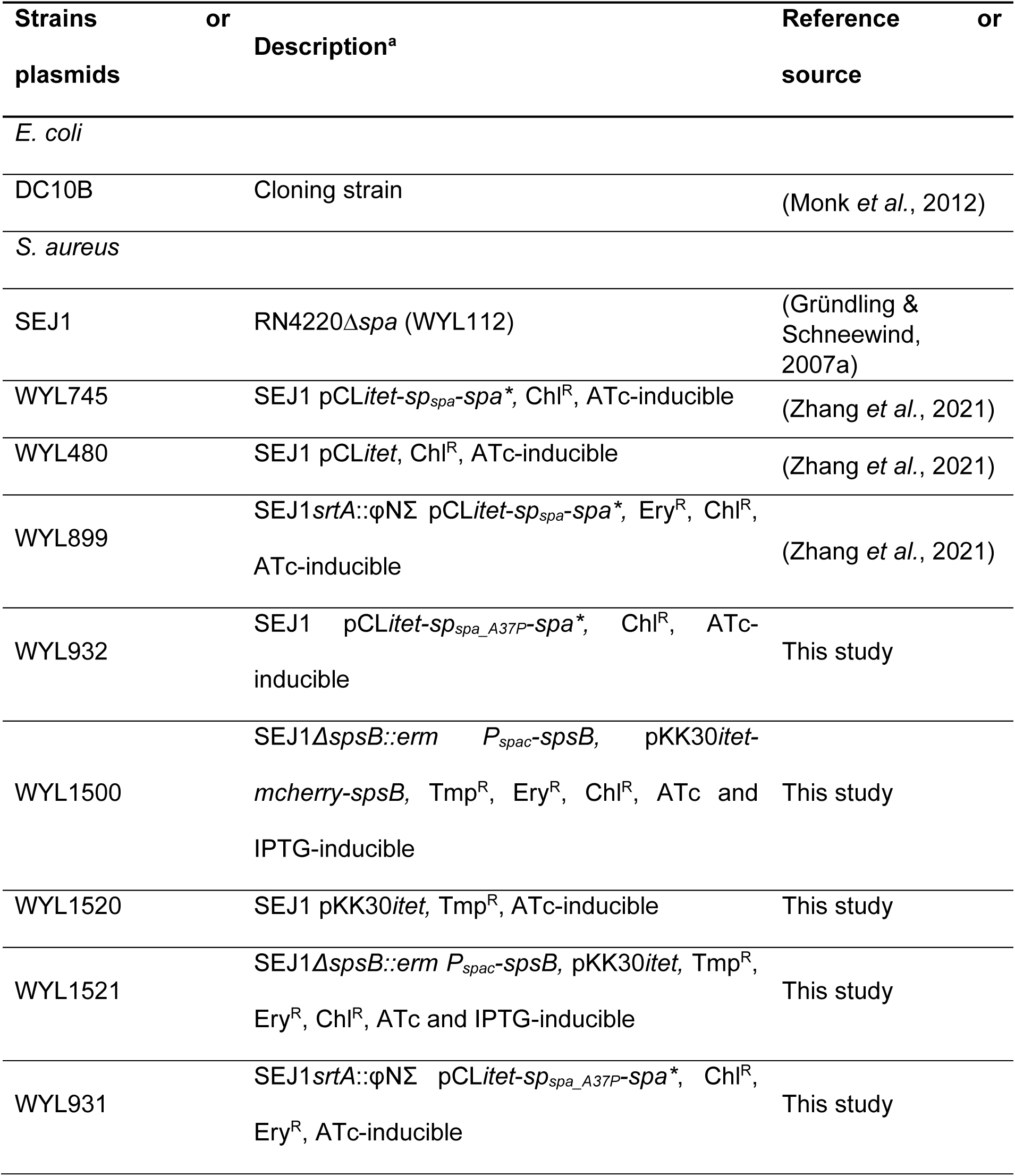
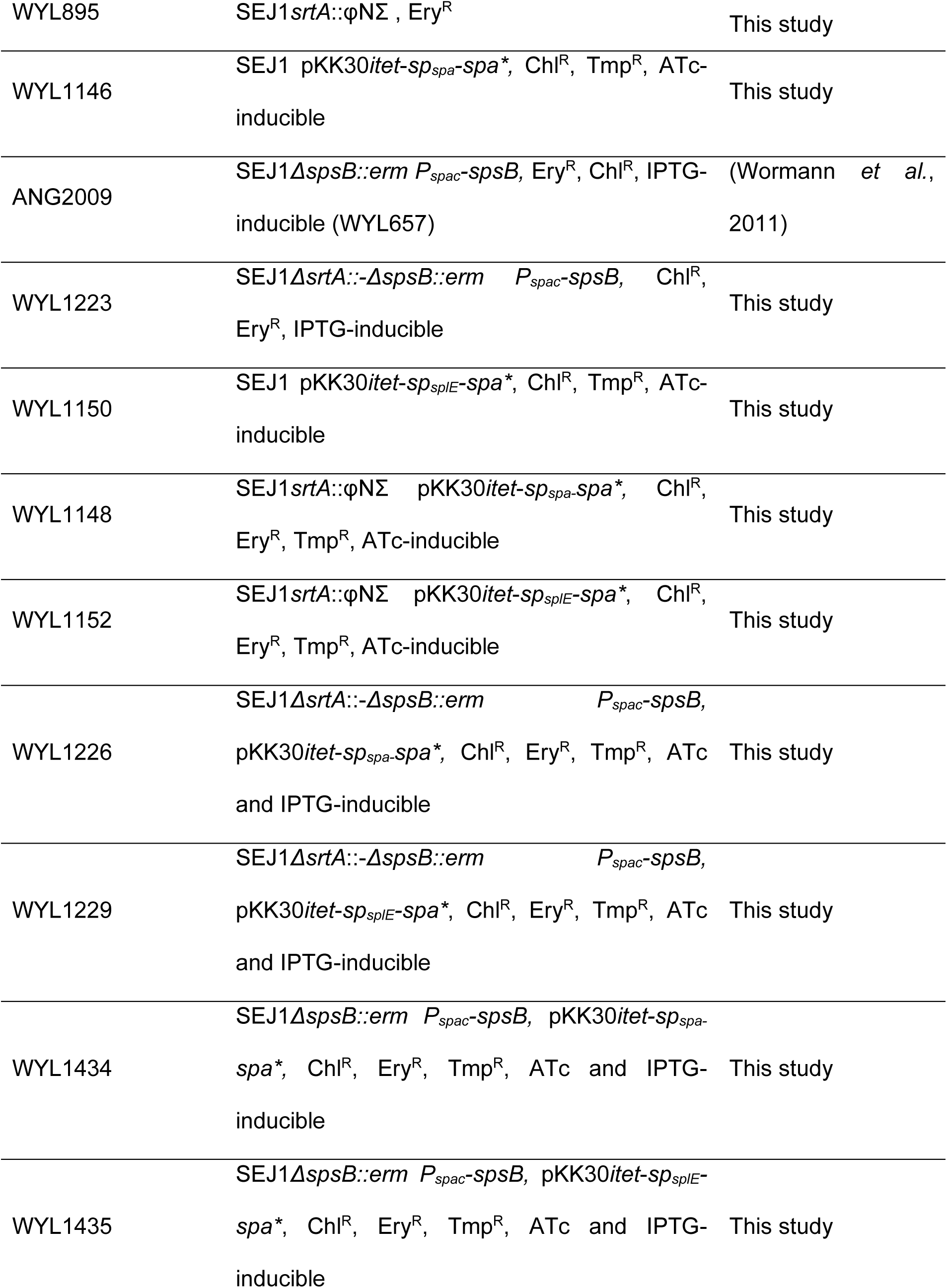
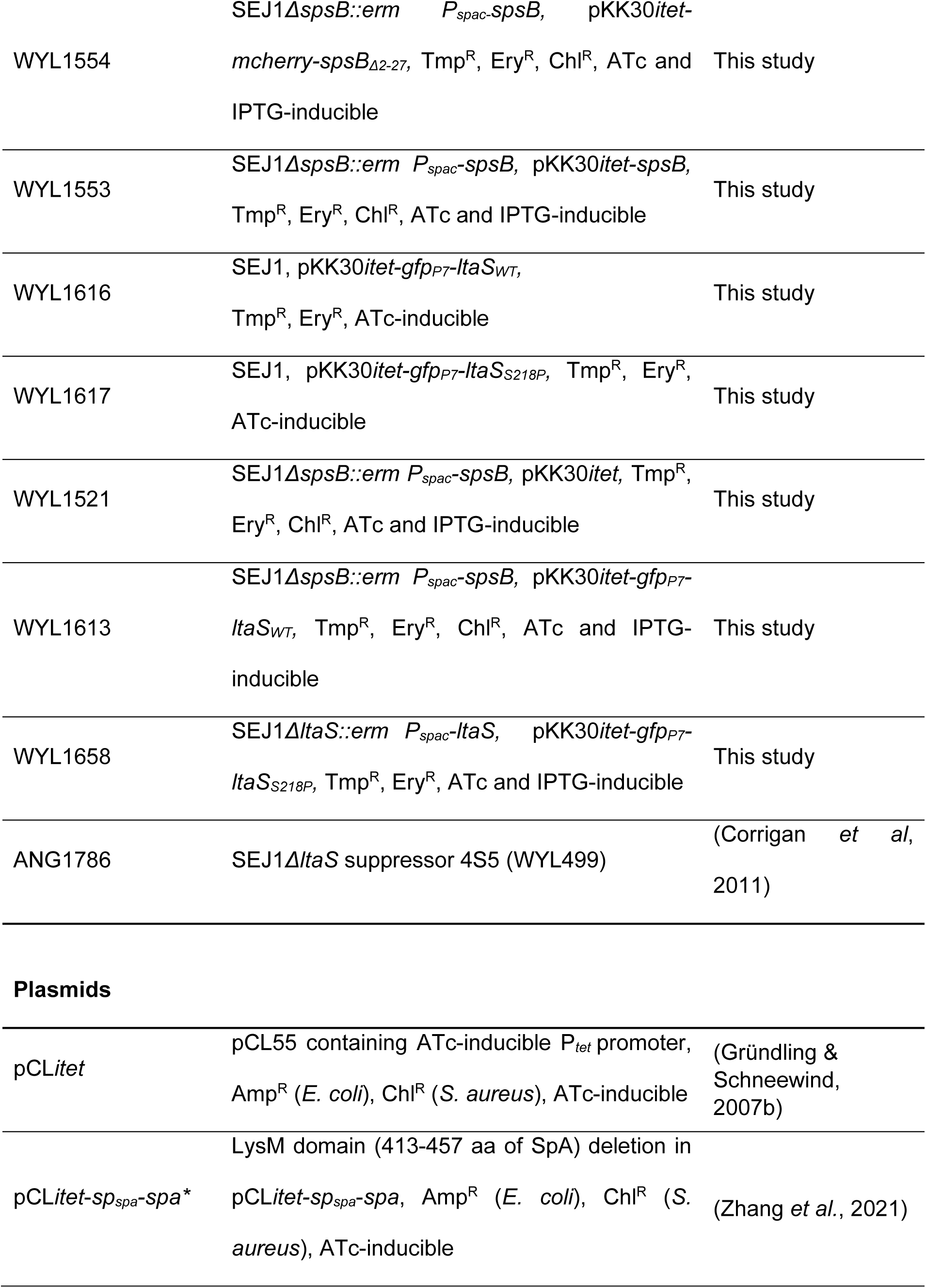
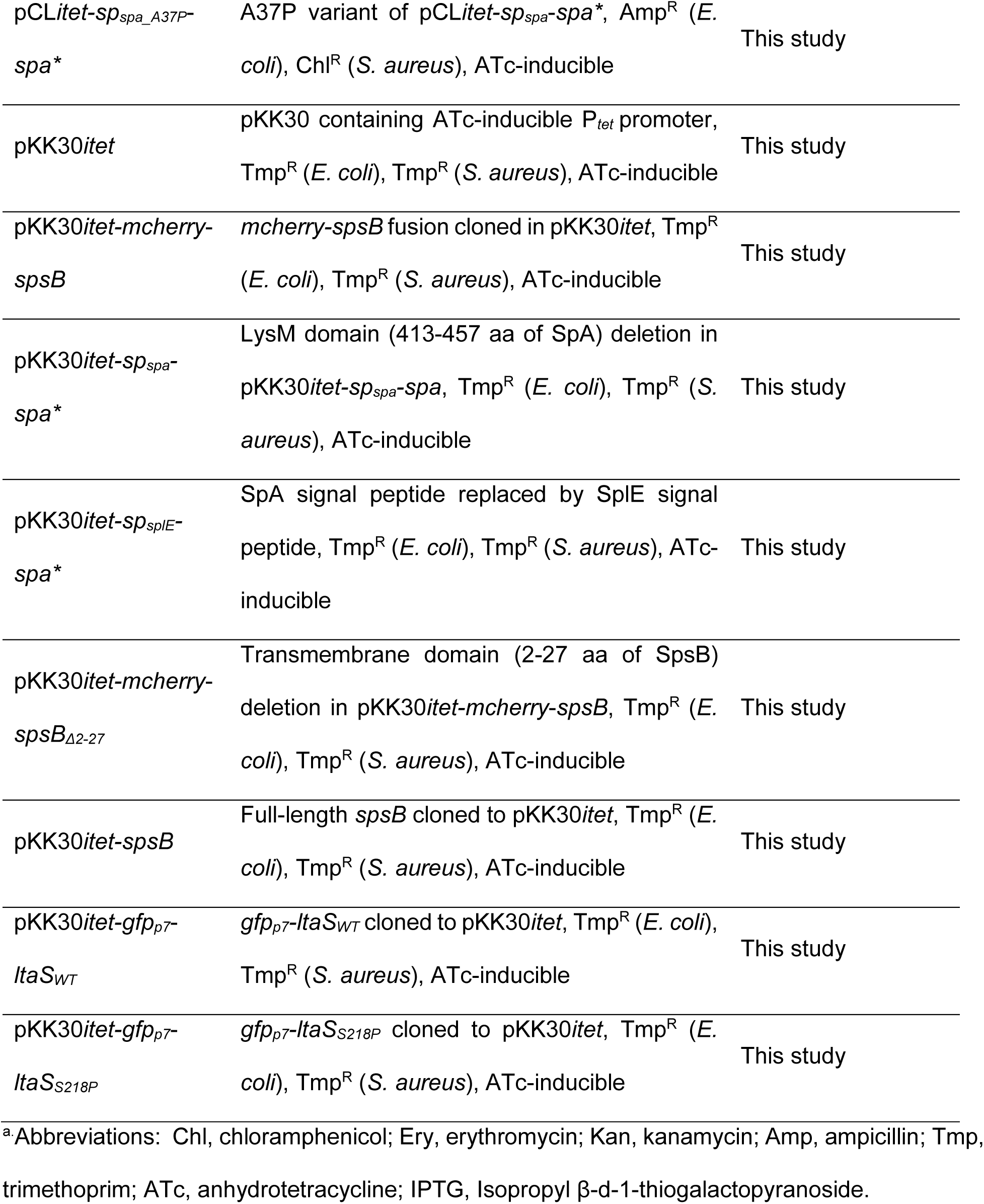
Strains and plasmids used in this study.

**Table S2.**
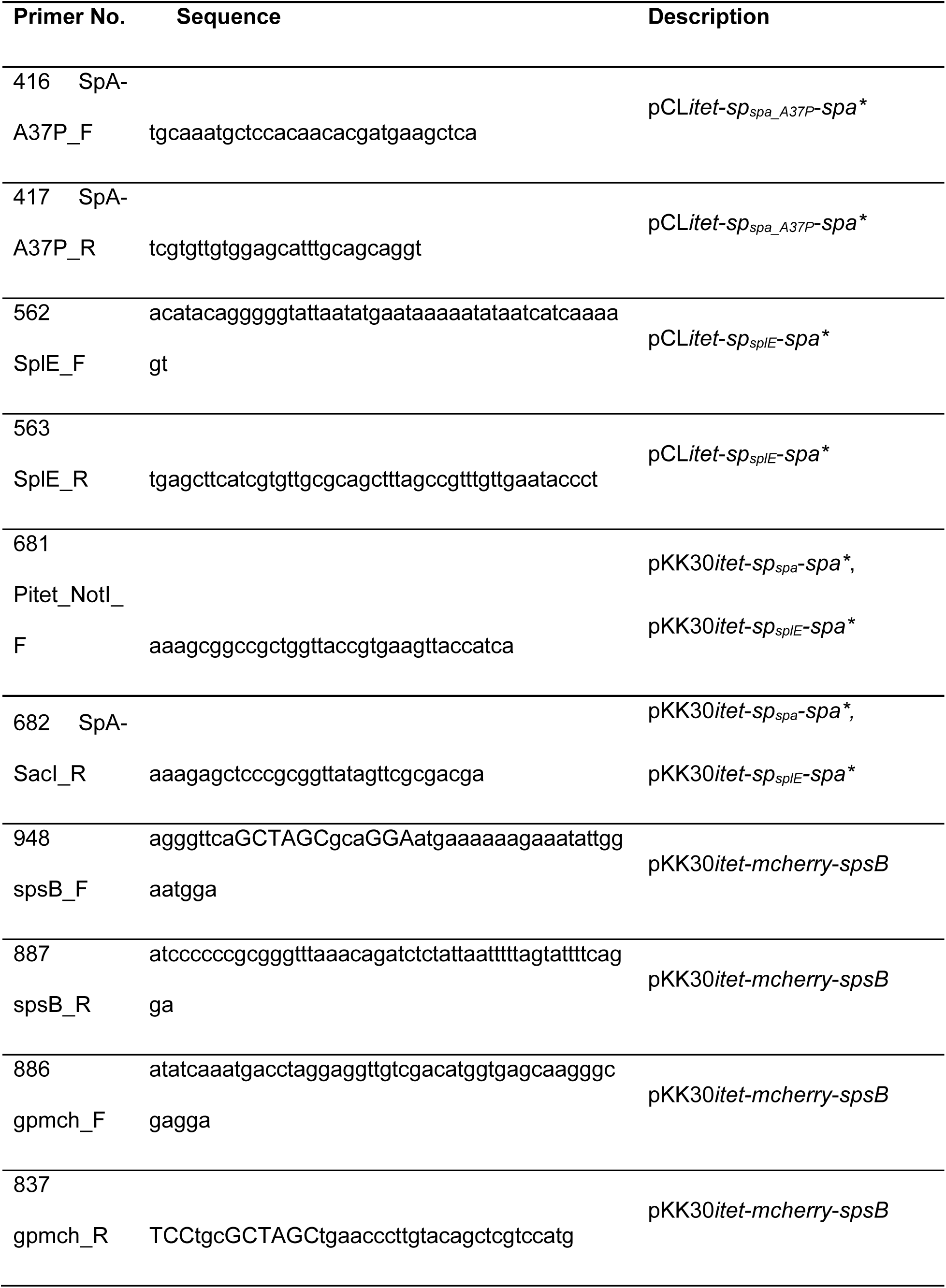
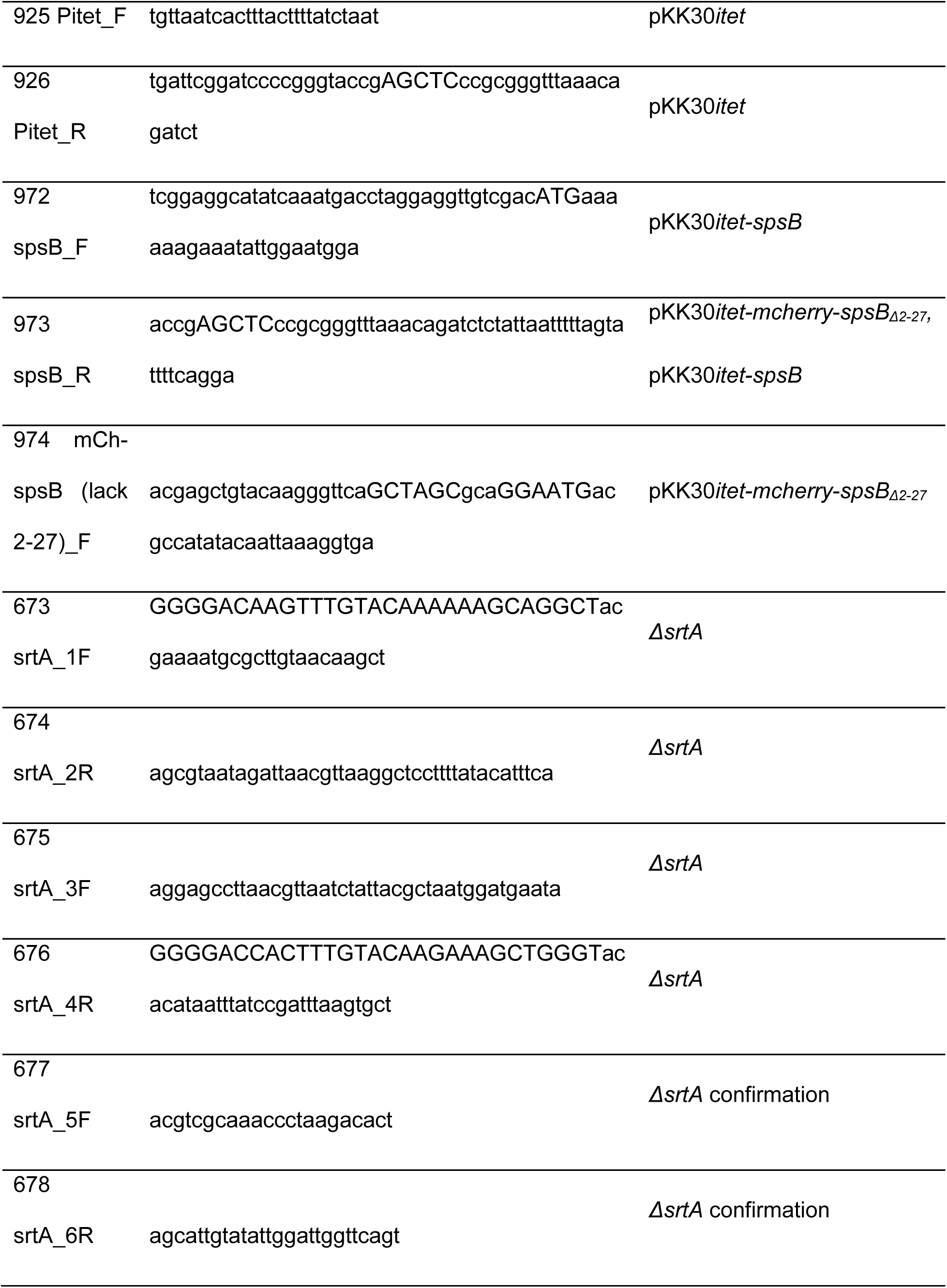
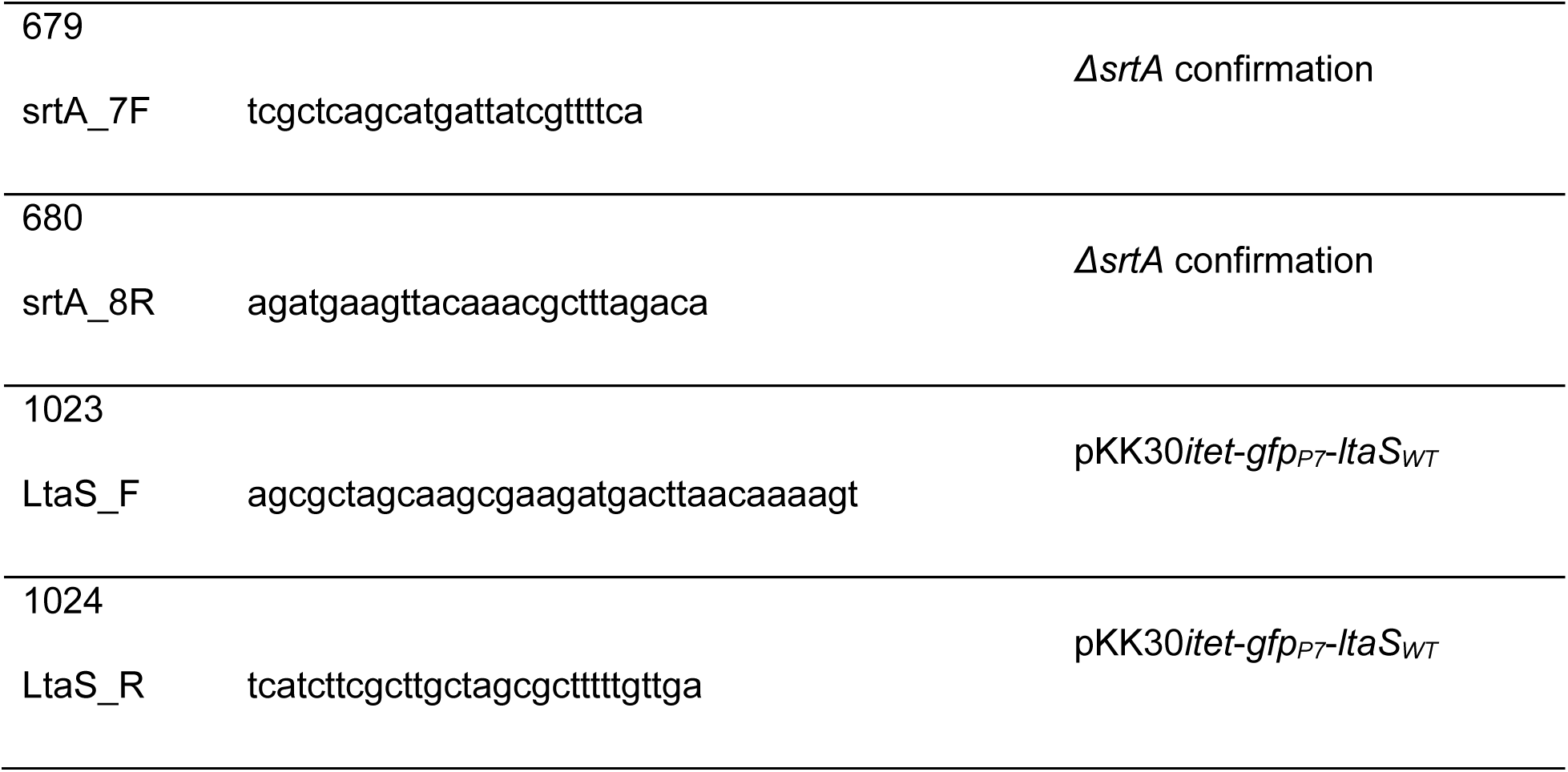
Primers used in this study.

